# NECTIN-4 PET FOR OPTIMIZING ENFORTUMAB VEDOTIN DOSE-RESPONSE IN UROTHELIAL CARCINOMA

**DOI:** 10.1101/2024.12.25.630315

**Authors:** Akhilesh Mishra, Ajay Kumar Sharma, Kuldeep Gupta, Dhanush R. Banka, Burles A. Johnson, Jeannie Hoffman-Censits, Peng Huang, David J. McConkey, Sridhar Nimmagadda

## Abstract

The optimization of dosing strategies is critical for maximizing efficacy and minimizing toxicity in drug development, particularly for drugs with narrow therapeutic windows such as antibody-drug conjugates (ADCs). This study demonstrates the utility of Nectin-4-targeted positron emission tomography (PET) imaging using [^68^Ga]AJ647 as a non-invasive tool for real-time assessment of target engagement in enfortumab vedotin (EV) therapy for urothelial carcinoma (UC). By leveraging the specificity of [^68^Ga]AJ647 for Nectin-4, we quantified dynamic changes in target engagement across preclinical models and established its correlation with therapeutic outcomes. PET imaging revealed dose-dependent variations in Nectin-4 engagement, with suboptimal EV doses resulting in incomplete Nectin-4 engagement and reduced tumor growth. Importantly, target engagement measured by PET emerged as a more reliable predictor of therapeutic efficacy than dose or baseline Nectin-4 expression alone. Receiver operating characteristic (ROC) analysis identified a target engagement threshold that is determinant of response, providing a quantitative benchmark for dose optimization. Furthermore, PET imaging measures provide a promising framework to account for key challenges in ADC development, including tumor heterogeneity, declining drug-to-antibody ratios over time, and limitations of systemic pharmacokinetic measurements to account for tumor-drug interactions. These findings underscore the transformative potential of integrating PET pharmacodynamic measures as early biomarkers to refine dosing strategies, improve patient outcomes, and accelerate the clinical translation of next-generation targeted therapeutics.

## INTRODUCTION

Determining the optimal dose is crucial in drug development, as it governs both clinical efficacy and patient safety. Suboptimal dosing can have significant consequences. Not only can it result in market withdrawal or the loss of therapeutic indications, but it also exposes patients to unnecessary toxicities, many of which may be severe or life-threatening, ultimately impacting their quality of life (1). Historically, dose-selection strategies have focused on identifying the maximum tolerated dose (MTD), an approach inherited from cytotoxic chemotherapy (2). While effective for older therapies, this method is increasingly viewed as inadequate for modern targeted therapies, such as antibody-drug conjugates (ADCs), which have a narrow therapeutic window and complex toxicity mechanisms, whereas immuno-oncology therapies generally offer a broader therapeutic window but involve complex response biology (1,3).

In response, initiatives such as the FDA’s Project Optimus, alongside industry efforts, have advocated for a paradigm shift in dose selection (4). These efforts emphasize the need for early and comprehensive evaluation of multiple doses during clinical trials, aiming to shift the focus from determining the highest tolerable dose to identifying an optimal balance between efficacy, safety, and tolerability (5). However, existing dose-selection paradigms continue to face challenges, particularly in accounting for the complexities of drug-target interactions within the tumor microenvironment (6). In this process, pharmacodynamic (PD) biomarkers remain an important and a challenging aspect of early phase drug development(7).

To address these limitations, we propose a dual approach that integrates early biomarker discovery with advanced imaging techniques. Imaging, particularly through methods such as positron emission tomography (PET), offers a complementary layer of data that enhances traditional pharmacokinetic (PK) and PD assessments. While PK/PD data provide insights into drug distribution and activity over time, imaging captures real-time, spatially-resolved measurements of drug distribution and drug-target interactions within the tumor and across the whole body(8). Combining these two streams of information offers a more precise understanding of dose-exposure-response relationships, enabling a shift from late-stage PK/PD modeling to a more comprehensive strategy that incorporates predictive biomarkers early in development.

In this context, we hypothesized that positron emission tomography (PET) imaging could significantly enhance our understanding of antibody-drug conjugate (ADC) pharmacology. ADCs hold promise for precise cancer targeting with minimal off-target effects, yet their optimization is complicated by interactions with the tumor microenvironment. Factors like high interstitial pressure and poor tumor vasculature can hinder ADC accumulation at the tumor site, complicating traditional plasma-based PK analyses (**Fig. 1**) (9). Additionally, conventional PK and PD assessments often fall short in capturing the complexities of drug-target interactions within tumors, especially given the variability in target expression across different cancers and the poorly understood factors in the tumor microenvironment that influence drug accumulation (6). PET offers real-time, non-invasive measurements of target expression in the whole body(8). By using custom-designed PET radiotracers that bind to the target with weaker affinities than the ADCs and designed for rapid clearance (10), we aimed to capture a clearer picture of accessible target levels at the tumor site and quantify the drug-tumor interactions. This real-time imaging approach complements the traditional PK/PD assessments and provides a more precise understanding of dose-exposure-response relationships.

**Figure 1.**
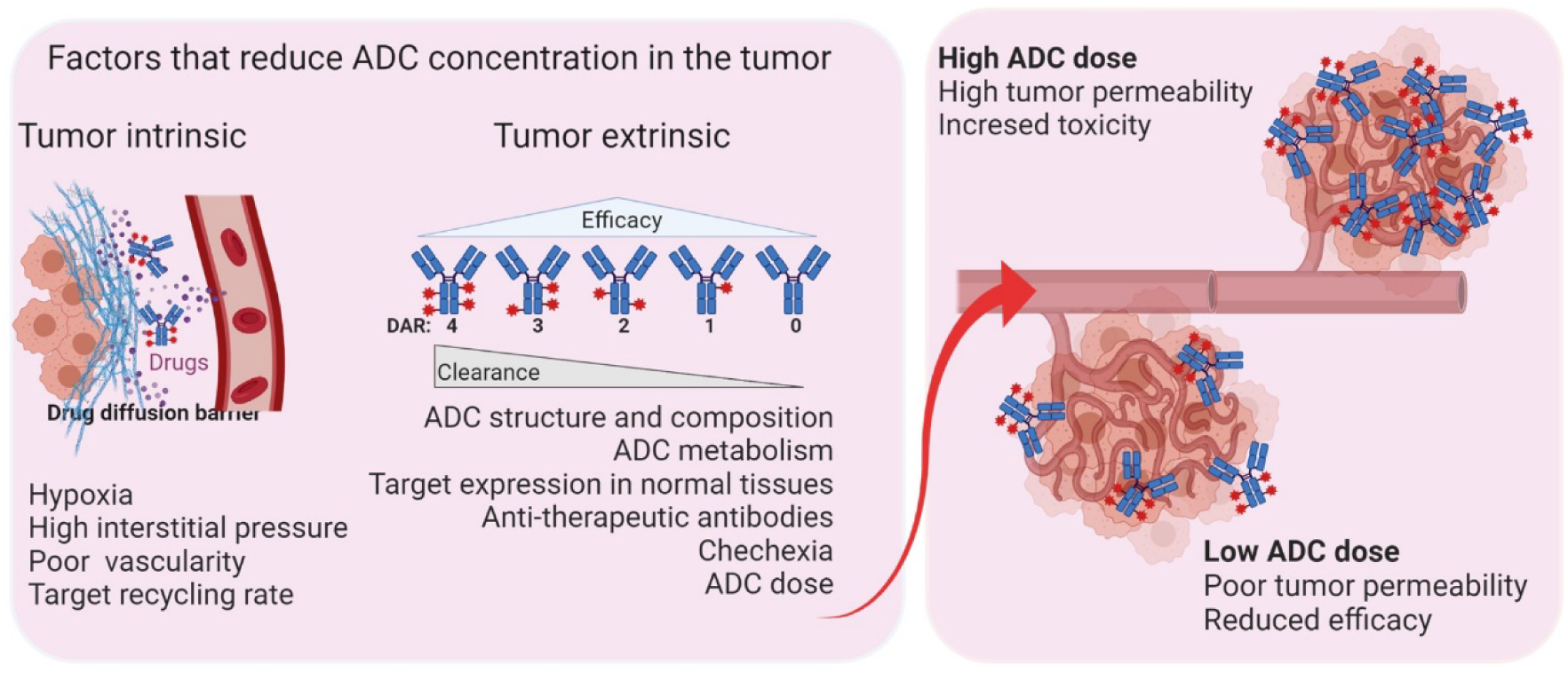
Schematic depicting intrinsic and extrinsic factors affecting antibody drug conjugate (ADC) concentration at the tumor. A tradeoff between toxicity and efficacy is critical for the long-term success of ADC therapy.

Focusing on enfortumab vedotin (EV), an FDA-approved ADC targeting nectin-4 (11), we sought to explore whether PET imaging measures of target engagement at the tumor, an early biomarker of drug-target interaction, could lead to more precise understanding of dose-exposure-response relationships. In the EV-101 study, although responses were observed at all dose levels of EV, the 1.25 mg/kg dose level was associated with greater tumor response, while lower response rates were observed at doses below 1.25 mg/kg (12). Adjustments in the dosage were more frequently necessary at the 1.25 mpk level, affecting over 32% of patients compared to those on lower dosages (12). These clinical findings indicate that understanding total nectin-4 levels and engagement dynamics is paramount to understanding the response to EV treatments. Nearly Although recent advances in nectin-4-targeted imaging agents have shown promise in quantifying target expression heterogeneity (13,14), they have not yet been applied to evaluate the pharmacodynamic effects of EV and to elucidate dose-exposure-response relationships. By leveraging PET to measure EV exposure within the tumor and correlate it with therapeutic response, we aimed to provide critical insights to optimize ADC dosing for improved efficacy and patient outcomes. Our results show that mapping target engagement dynamically and early in the treatment process could potentially lead to more effective and safer balance between efficacy and toxicity in ADC therapies and other targeted treatments.

## RESULTS

### Synthesis and characterization of [^68^Ga]AJ647 as Nectin-4 binding moiety

AJ647, derived from AJ632—a 15-amino-acid bicyclic peptide with low nanomolar affinity to Nectin-4 (15)— was further optimized to enhance its pharmacokinetic properties. A PEG linker was attached to the N-terminal cysteine residue of AJ632 (**Fig. 2A**). The free terminal amine was then conjugated to a NOTA chelator, enabling the successful radiolabeling with gallium-68 to produce [^68^Ga]AJ647 and its non-radioactive analog [^nat^Ga]AJ647. The radiolabeling process, performed as previously described (16), yielded [^68^Ga]AJ647 with an 71.2±9.4% decay-corrected yield and 97±0.8% purity (n=20), achieving high molar specific activity (**Fig. S1-S6**). The stability of [^68^Ga]AJ647 was confirmed for up to 3 hours in its final formulation. Additionally, the compound exhibited a logD value of -2.06±0.17, indicating high solubility in aqueous media (**Fig. S7**). Surface plasmon resonance studies demonstrated that [^nat^Ga]AJ647 bound to human and mouse Nectin-4 with affinities of 15.0±10.1 nM and 14.1±3.4 nM, respectively (**Fig. 2B-C**).

**Figure 2.**
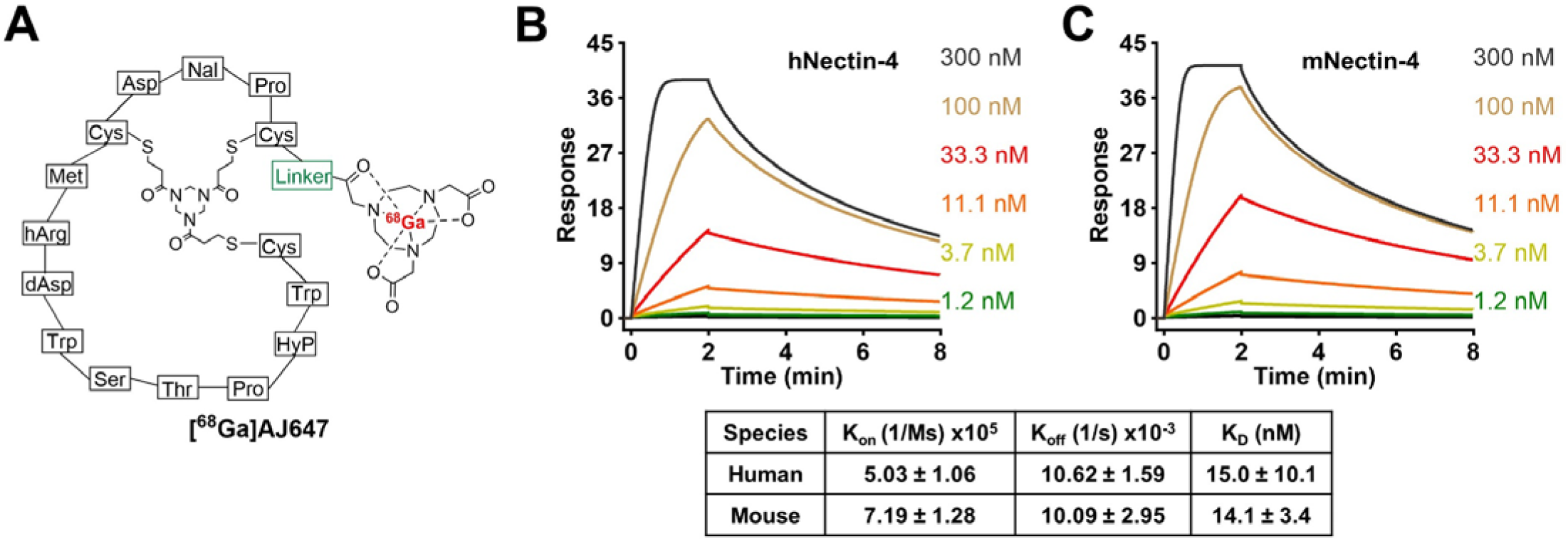
Structural design and in vitro affinity assessment of AJ647. **A)** Molecular structure of AJ647, a bicyclic peptide having NOTA as a bifunctional chelator for ^68^Ga-labeling. **B)** and **C)** Surface Plasmon Resonance (SPR) analysis showing the binding affinity of AJ647 towards Nectin-4, assessed using recombinant human and mouse nectin-4 proteins respectively. Data is presented as mean±SD (n=3).

To assess the binding specificity of [^68^Ga]AJ647 to Nectin-4, we selected several UC cell lines, including T24, SCaBER, RT112, UC9, UC14 and HT1376. Upon incubation with [^68^Ga]AJ647, HT1376 cells demonstrated the highest uptake (6.5±0.52), followed by UC14 (3.0±0.42), UC9 (2.2±0.48), RT112 (1.9±0.18), SCaBER (1.6±0.13), and T24 (0.2±0.11) (**Fig. 3A**). Flow cytometry analysis corroborated these results, showing a consistent pattern of Nectin-4 expression across these cell lines, with expression levels ranked as follows: T24 < SCaBER < RT112 < UC9 < UC14 < HT1376 (**Fig. 3B**). Furthermore, the quantification of cell surface Nectin-4 density using PE-labeled beads revealed a strong correlation between receptor density and [^68^Ga]AJ647 uptake (**Fig. 3C**, R^2^=0.97). Collectively, these findings demonstrate the high in vitro selectivity of [^68^Ga]AJ647 for Nectin-4.

**Figure 3.**
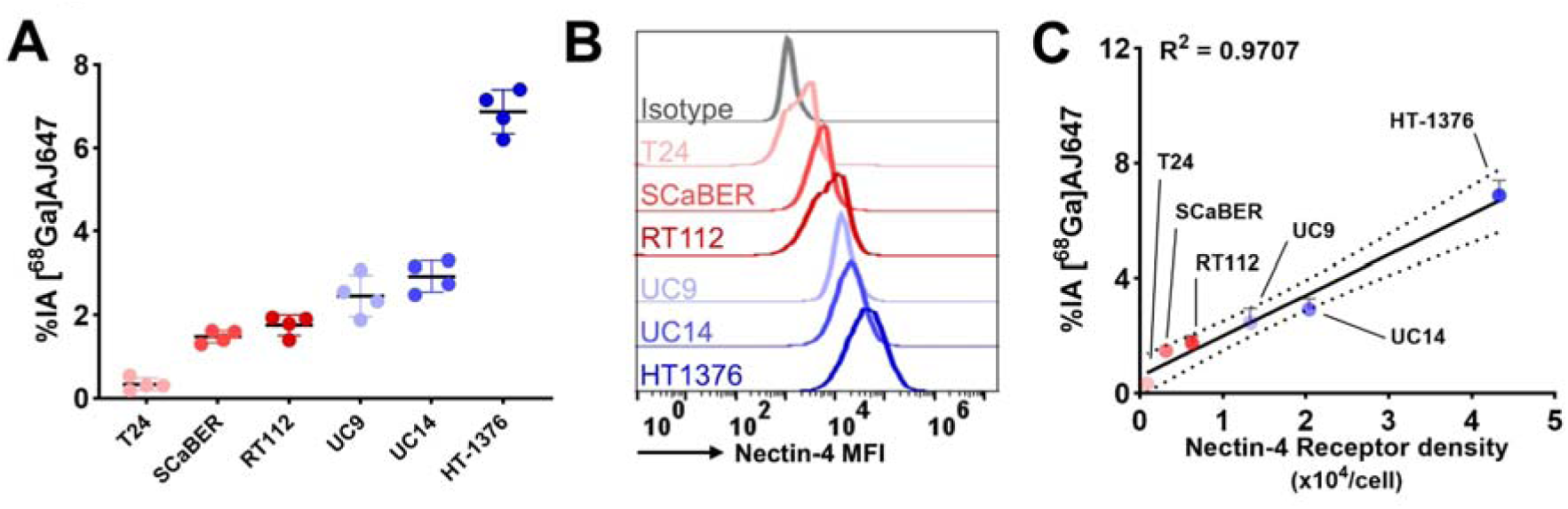
In vitro specificity of [^68^Ga]AJ647 for Nectin-4 in bladder cancer cell lines. **A)** [^68^Ga]AJ647 binding (percent incubated activity, %IA) to various bladder cancer cell lines. Cells were incubated with ∼37 kBq (∼1 µCi) [^68^Ga]AJ647 at 4°C for 1 hour. Specificity was confirmed by pre-incubating with 2 μM of non-radioactive AJ647 (blocking dose), which significantly reduced radiotracer uptake, indicating that [^68^Ga]AJ647 targets Nectin-4. **B)** Flow cytometry analysis of Nectin-4 receptor expression across different bladder cancer cell lines, providing insight into the variability in receptor density among the cell lines. **C)** Correlation between in vitro [^68^Ga]AJ647 uptake and Nectin-4 receptor density, demonstrating that the uptake of [^68^Ga]AJ647 is dependent on cell surface density of Nectin-4 receptors. Data in figure A and C are represented as mean ± SD. Pearson correlation used in C.

### Pharmacokinetics of [^68^Ga]AJ647

Given that HT1376 exhibited the highest Nectin-4 expression, we conducted a pharmacological evaluation of [^68^Ga]AJ647 in HT1376 xenografts using immunocompromised NOD SCID γ^NULL^(NSG) mice. Dynamic PET-MR imaging was performed for up to 90 minutes post-injection. PET images revealed significant radiotracer accumulation in HT1376 tumors as early as 15 minutes post-injection (**Fig. 4A**). By 60 minutes, the tumor contrast had further improved due to background radiotracer clearance, as evidenced by the percent injected activity per cubic centimeter (%IA/cc) values derived from [^68^Ga]AJ647 PET (**Fig. 4B**). The specificity of [^68^Ga]AJ647 for Nectin-4 was confirmed by the reduction in tracer uptake in HT1376 xenografts following administration of 1 mg/kg of non-radioactive AJ647 (**Fig. S8**). Notably, [^68^Ga]AJ647 uptake was also observed in the skin and salivary glands, consistent with the expression of Nectin-4 in these tissues (**Fig. S8**). In contrast, minimal uptake was detected in non–Nectin-4-expressing tissues such as muscle. Among normal tissues, the kidneys and bladder showed the highest radiotracer accumulation, consistent with the renal clearance mechanism typical for low-molecular-weight peptides (**Fig. 4B**).

**Figure 4.**
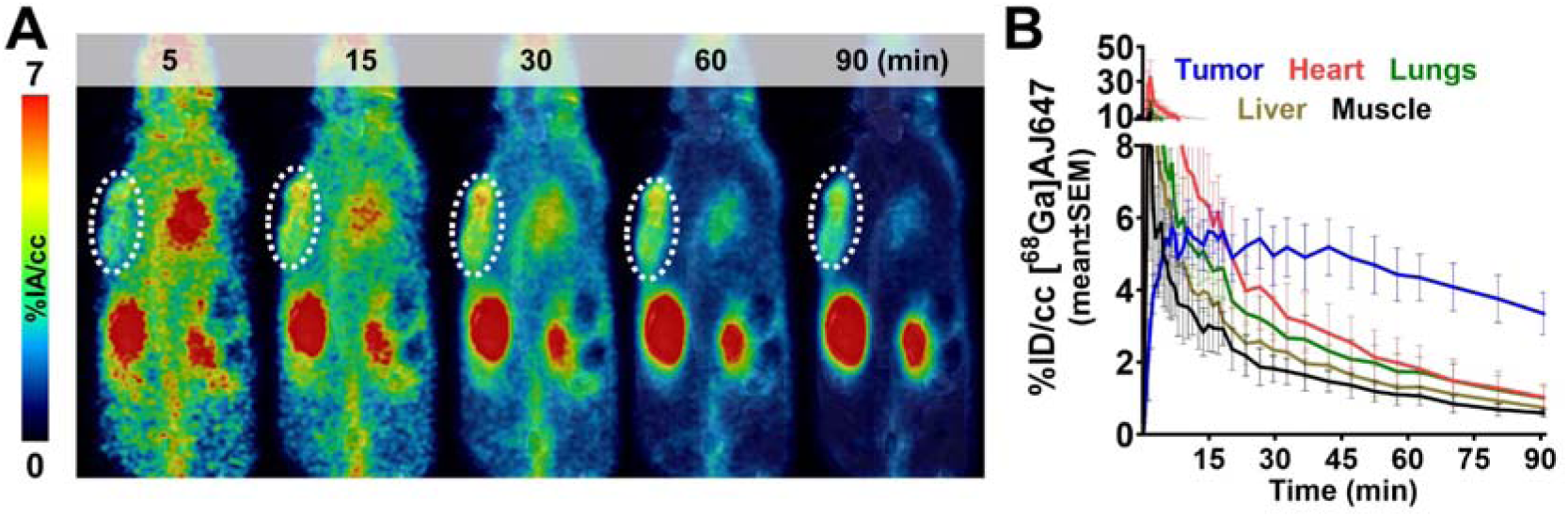
Pharmacokinetics of [^68^Ga]AJ647 in NSG mice bearing HT1376 tumors. **A)** Dynamic PET imaging of NSG mice injected with ∼9.25 MBq (∼250 µCi) of [^68^Ga]AJ647 and images were acquired continuously from -1 to 90 minutes. The coronal section of fused PET/MR images at selected time points shows the accumulation and retention of [^68^Ga]AJ647 in the tumor (indicated by a white circle) and radiotracer’s washout from background tissues **B)** Time-activity curves of [^68^Ga]AJ647 uptake in the tumor, heart, lungs, liver and muscle derived from dynamic scans from -1 to 90 min, showing differential uptake and clearance of [^68^Ga]AJ647 in these tissues over time, highlighting its preferential retention in the tumor compared to other organs. Data in figures **B** is represented as mean±SEM (n =3).

To corroborate the PET imaging results, ex vivo biodistribution studies were performed **(Table 1)**. [^68^Ga]AJ647 uptake in Nectin-4-expressing tumors reached a maximum of 3.75±0.86 %IA/g at 15 minutes, with similar retention observed at 60 minutes (3.72±0.36). Blood pool activity demonstrated efficient clearance, with over 90% of the activity cleared by 60 minutes. Comparable clearance patterns were observed in all other non-specific tissues. Consequently, high tumor-to-muscle (10.07±2.19) and tumor-to-blood (4.60±1.07) ratios were achieved at 60 minutes. Based on these findings, we determined that the 60-minute time point provided optimal contrast and aligned well with standard PET clinical workflows, making it the preferred time point for subsequent experiments.

**Table 1.** Pharmacokinetics of [^68^Ga]AJ647 in mice bearing HT1376 tumors. Mice bearing HT1376 tumors were injected with ∼1.85 MBq (∼50 µCi) of [^68^Ga]AJ647 in 200 µL saline containing 5% ethanol. Mice were sacrificed at specified time points post-injection, and tissues were harvested for analysis (n=5 per group).

| %IA/g or ratio | Time (min) |  |  |  |  |
| --- | --- | --- | --- | --- | --- |
|  | 2 | 15 | 60 | 120 | 180 |
| Blood | 35.75±3.64 | 2.94±0.68 | 0.84±0.22 | 0.28±0.06 | 0.32±0.09 |
| Skin | 1.13±0.16 | 2.74±0.50 | 1.45±0.33 | 0.82±0.22 | 0.66±0.12 |
| Muscle | 0.96±0.19 | 0.93±0.36 | 0.38±0.06 | 0.20±0.11 | 0.18±0.05 |
| Femur | 1.88±0.29 | 1.30±0.05 | 0.57±0.18 | 0.27±0.09 | 0.34±0.09 |
| Salivary glands | 6.30±1.24 | 4.74±0.94 | 1.48±0.18 | 0.67±0.09 | 0.69±0.07 |
| Tumor | 0.72±0.31 | 3.75±0.86 | 3.72±0.36 | 2.06±0.49 | 1.39±0.32 |
| Lung | 24.18±7.05 | 3.66±0.50 | 1.66±0.25 | 0.81±0.28 | 0.47±0.20 |
| Thymus | 15.69±5.42 | 2.38±0.70 | 0.71±0.30 | 0.67±0.21 | 0.52±0.23 |
| Heart | 11.34±0.51 | 1.15±0.34 | 0.43±0.07 | 0.19±0.01 | 0.22±0.01 |
| Liver | 6.45±1.39 | 1.19±0.30 | 0.45±0.09 | 0.21±0.07 | 0.23±0.02 |
| Stomach | 0.34±0.11 | 0.49±0.36 | 0.27±0.01 | 0.13±0.04 | 0.16±0.04 |
| Small Intestine | 0.85±0.23 | 0.68±0.23 | 0.52±0.17 | 0.16±0.02 | 0.28±0.02 |
| Pancreas | 2.39±0.72 | 2.50±0.38 | 0.93±0.03 | 0.30±0.06 | 0.42±0.02 |
| Spleen | 1.04±0.23 | 4.32±1.31 | 1.75±0.21 | 0.84±0.17 | 0.83±0.07 |
| Large Intestine | 0.25±0.16 | 0.41±0.11 | 0.25±0.13 | 0.18±0.06 | 0.28±0.13 |
| Adrenals | 4.81±2.58 | 2.60±1.10 | 4.45±2.03 | 3.11±0.21 | 3.01±0.27 |
| Kidney | 4.32±1.31 | 20.43±3.7 | 30.82±4.37 | 17.33±12.86 | 24.82±8.74 |
| Bladder | 1.72±0.24 | 24.13±13.51 | 39.52±21.31 | 22.60±6.26 | 58.04±24.76 |
| Testicles | 0.79±0.07 | 0.86±0.12 | 0.71±0.24 | 0.52±0.17 | 0.93±0.50 |
| Brain | 0.54±0.15 | 0.10±0.02 | 0.06±0.01 | 0.06±0.02 | 0.07±0.01 |
| Tumor/Blood | 0.02±0.01 | 1.31±0.41 | 4.60±1.07 | 7.27±0.76 | 4.58±1.33 |
| Tumor/Muscle | 0.79±0.45 | 4.23±0.94 | 10.07±2.19 | 11.26±3.03 | 8.20±4.02 |

### Evaluation of [^68^Ga]AJ647 specificity in UC xenografts

To confirm the in vivo specificity of [^68^Ga]AJ647 for detecting varying levels of Nectin-4, we conducted studies using xenografts derived from the six UC cell lines previously use in our in vitro experiments. Consistent with our in vitro findings, the highest uptake of [^68^Ga]AJ647 was observed in HT1376 tumors, followed by UC14, UC9, RT112, SCaBER, with the lowest uptake in T24 tumors (**Fig. 5A**). These findings were corroborated by immunohistochemical (IHC) staining of the same xenografts for Nectin-4 expression, which closely mirrored PET imaging results (**Fig. 5B**).

**Figure 5.**
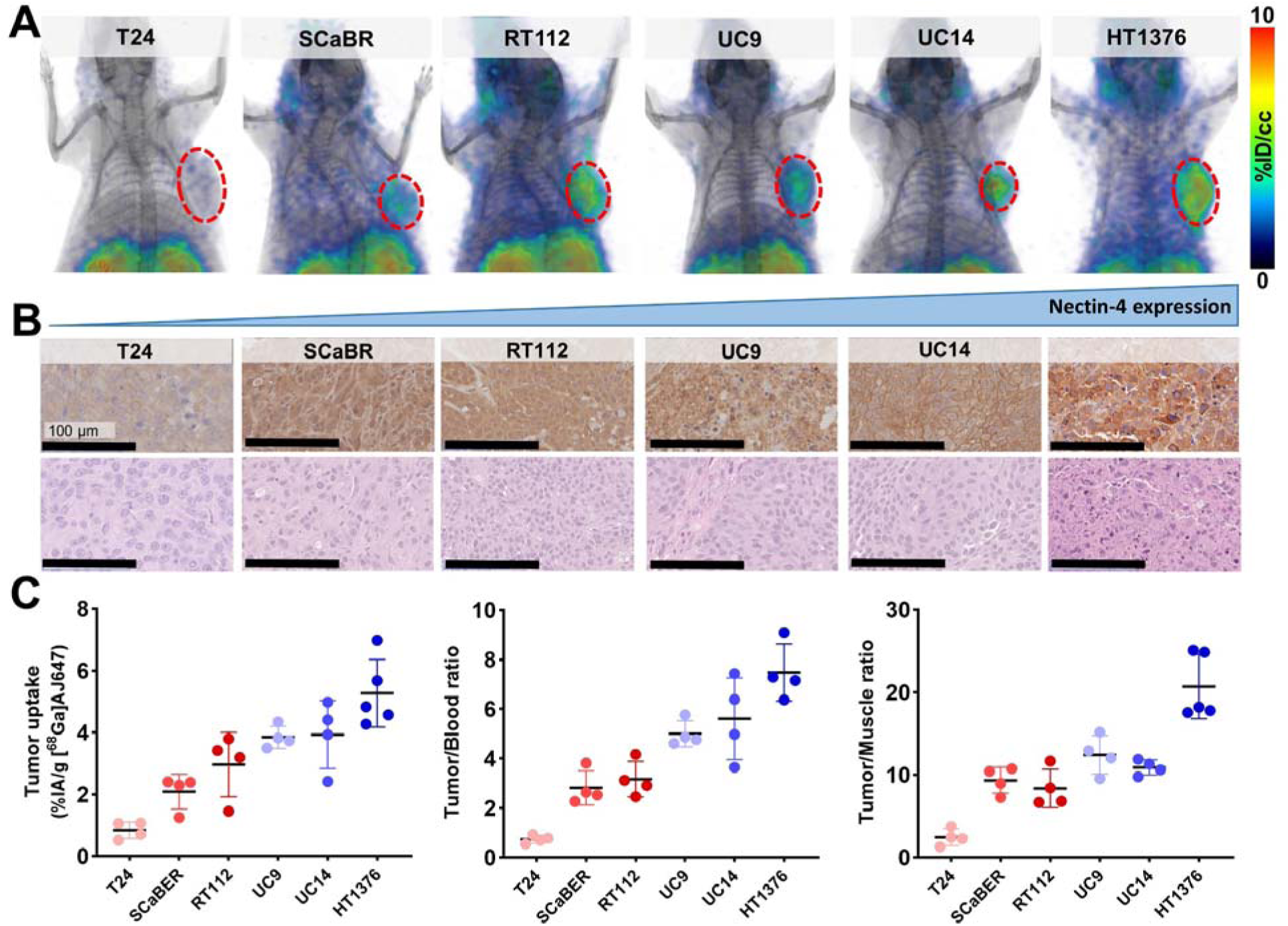
In vivo specificity of [^68^Ga]AJ647 for Nectin-4 in UC tumor xenografts. **A)** Whole-body static PET-CT images acquired 60 minutes after intravenous injection of approximately 7.4 MBq (∼200 µCi) [^68^Ga]AJ647 in NSG mice bearing UC tumor xenografts. 3D volume-rendered PET images reveal significantly higher radiotracer accumulation in the tumors (highlighted by a red circle) compared to background tissues, demonstrating the specificity of [^68^Ga]AJ647 for Nectin-4. **B)** IHC staining of Nectin-4 in corresponding tumor sections. The staining reveals variable Nectin-4 receptor densities across the tumor xenografts. **C)** Tumor uptake, tumor-to-blood and tumor-to-muscle ratios derived from ex vivo biodistribution studies. NSG mice bearing UC xenografts were injected with approximately 740 kBq (∼20 µCi) [^68^Ga]AJ647 and sacrificed 60 minutes post-injection. Ex vivo biodistribution data is showing the selective accumulation of [^68^Ga]AJ647 in tumors relative to blood and muscle. Data in **C** are represented as mean ± SD (n=4-5).

To validate these imaging findings, we performed ex vivo biodistribution studies. The results confirmed that tumor uptake of [^68^Ga]AJ647 was consistent with both the in vitro data and the PET imaging outcomes. Specifically, the highest uptake was observed in HT1376 xenografts (%IA/g: 5.27±1.09), followed by UC14 (3.93±1.10), UC9 (3.85±0.36), RT112 (2.96±1.04), SCaBER (2.09±0.56), and the lowest in T24 xenografts (0.84±0.27), which corresponded well with their respective Nectin-4 expression levels (**Fig. 5C, left**). Additionally, tumor-to-blood (**Fig. 5C**, **middle**) and tumor-to-muscle (**Fig. 5C**, **right**) ratios mirrored these trends, further supporting the specificity of [^68^Ga]AJ647 for Nectin-4. Background tissues displayed no significant differences across the tumor models (**Fig. S9**).

Collectively, these results demonstrate that [^68^Ga]AJ647 is highly specific to Nectin-4 and effectively detects its varying expression levels in tumors in vivo.

### [^68^Ga]AJ647-PET could be used to quantify the Nectin-4 engagement of EV in vitro and in vivo

Previously, we have demonstrated that low molecular weight peptide-based imaging agents that overlap in binding mode with the therapeutic antibodies could be used to quantify the target engagement of those therapeutics by measuring unoccupied (accessible) target levels(10,17). To investigate if [^68^Ga]AJ647 could be used similarly to investigate the target engagement of EV we performed in vitro competition assays (**Fig. 6A**). Initially, HT1376 cells were incubated with [^68^Ga]AJ647 in the presence of increasing concentrations of EV. The results revealed a dose-dependent reduction in [^68^Ga]AJ647 binding, with an IC_50_ of 2.7 nM (**Fig. 6B**). We then performed a reverse competition assay using fluorescently labeled EV (EV-FITC) as a probe while varying concentrations of non-radioactive AJ647 were introduced. This led to a similar dose-dependent reduction in EV-FITC binding, with an IC_50_ of 35 nM (**Fig. 6C**). To validate that these observations are applicable across variable Nectin-4 expression, we treated HT1376, SCaBER and T24 cells that exhibit variable Nectin-4 expression levels with EV for 15 min (60 nM) and incubated with [^68^Ga]AJ647. We observed that in all the cell lines tested, presence of EV significantly reduced [^68^Ga]AJ647 binding in Nectin-4 positive HT1376 and ScaBER cells(P<0.0001) but not in Nectin-4 negative T24 cells (**Fig. 6D**). These findings confirm that [^68^Ga]AJ647 binds to the same epitope on Nectin-4 as that of EV. Notably, the weaker binding affinity (higher IC_50_) of AJ647 for Nectin-4, compared to EV, suggests that [^68^Ga]AJ647, used at pM-range doses typically, is unlikely to disrupt EV-Nectin-4 interactions and binds only to accessible Nectin-4.

**Figure 6.**
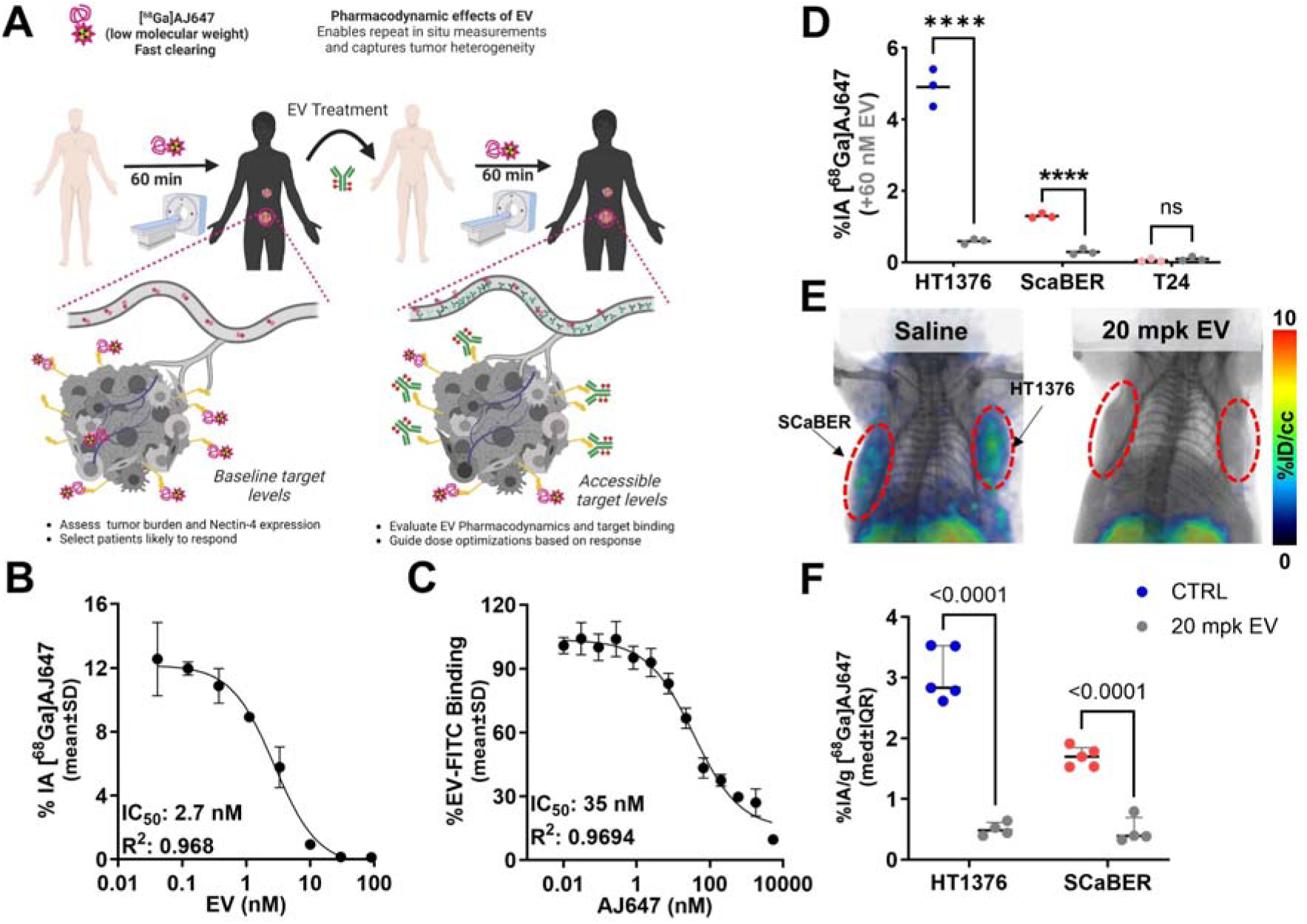
Competition binding assays of [^68^Ga]AJ647 and EV-FITC with Nectin-4 in HT1376 cell line. **A)** Schematic representation of competition assays between [^68^Ga]AJ647 and EV for binding to Nectin-4. On the left, the schematic illustrates the impact of varying EV concentrations on Nectin-4 levels, with accessible Nectin-4 quantified using [^68^Ga]AJ647. On the right, the schematic depicts how varying concentrations of AJ647 affect Nectin-4 levels, with accessible Nectin-4 determined by EV-FITC binding. **B)** In vitro binding of [^68^Ga]AJ647 to HT1376 cells with varying concentrations of EV. The data show concentration-dependent inhibition of [^68^Ga]AJ647 binding, with an IC_50_ of 2.7 nM, indicating the competitive nature of EV in blocking [^68^Ga]AJ647 binding to Nectin-4. **C)** In vitro binding of EV-FITC to HT1376 cells, assessed by flow cytometry, with varying concentrations of AJ647. The results demonstrate concentration-dependent inhibition of EV-FITC binding, with an IC_50_ of 35 nM, highlighting that AJ647 competes with EV-FITC for Nectin-4 binding. data represented as mean±SD (n=3-4). **D)** In vitro binding of [^68^Ga]AJ647 to HT1376, SCaBER and T24 cells with and without 60 nM EV. **E)** In vivo PET-CT imaging in mice harboring SCaBER (left) and HT1376 (right) tumors pre- and post-20 mpk EV treatment (n=2). PET-CT was acquired 60-minutes after 7.4 MBq (∼200 µCi) [^68^Ga]AJ647 injection. **E)** In vivo PET-CT imaging in mice harboring SCaBER (left) and HT1376 (right) tumors at saturating dose of 20 mpk or saline (n=2). PET-CT was acquired at 60-minutes after 7.4 MBq (∼200 µCi) [^68^Ga]AJ647 injection. **F)** Ex vivo biodistribution in mice harboring SCaBER (left) and HT1376 (right) tumors treated with 20 mpk or saline control (n=5). Mice were sacrificed at 60-minutes after 1.85 MBq (∼50 µCi) [^68^Ga]AJ647 injection.

To assess Nectin-4 engagement by EV in vivo in a non-invasive manner, we utilized two UC xenograft models, which exhibit varying levels of Nectin-4 expression. These models were selected to investigate the ability of EV to target Nectin-4 across different tumor profiles, as Nectin-4 expression is heterogeneous in UC and serves as a key therapeutic target(18). NSG mice bearing HT1376 and SCaBER cell-derived xenografts, which represent high and moderate Nectin-4 expression levels respectively, were treated with a single dose of EV (15 mg/kg) administered intravenously 24 hours prior to the injection of [^68^Ga]AJ647. PET images were acquired 1 hour post-injection of [^68^Ga]AJ647 in HT1376 model (**Fig. 6E**). Mice treated with EV exhibited a significant reduction in [^68^Ga]AJ647 uptake in tumors compared to saline-treated controls, indicating successful Nectin-4 engagement by EV.

The PET imaging data were corroborated by ex vivo biodistribution analysis, which showed a significant reduction in [^68^Ga]AJ647 uptake in EV-treated tumors compared to controls: 84% reduction in HT1376 tumors (P < 0.0001) and 72% reduction in SCaBER tumors (P < 0.001) (**Fig. 6F**). These results demonstrate that [^68^Ga]AJ647 PET imaging can be effectively used to quantify in vivo Nectin-4 target engagement by EV, offering a valuable tool for monitoring drug-target interactions in tumors with varying Nectin-4 expression.

The decrease in radiotracer accumulation was more pronounced in HT1376 tumors, correlating with their higher baseline Nectin-4 expression. These findings suggest a saturation of Nectin-4 by EV, particularly in tumors with greater receptor availability.

### Nectin-4 PET Quantifies Dose-Exposure Relationship of EV at the Tumor

ADCs, while designed to achieve a broad therapeutic window, often exhibit a relatively narrow margin compared to most monoclonal antibodies (1). This narrower therapeutic window frequently leads to dose reductions or early discontinuation of treatment due to toxicity concerns (1). A better understanding of dose-exposure dynamics, especially real-time monitoring of target engagement at the tumor, could enable more precise adjustments to dosing regimens and potentially expand the therapeutic window of ADCs.

In this study, we investigated whether Nectin-4 PET imaging could provide a quantitative assessment of changes in accessible Nectin-4 levels over time and track target engagement at the tumor in response to varying doses of EV. To evaluate this, we administered EV at doses of 6, 9, or 15 mg/kg, which correspond to FDA-approved human doses of 0.5, 0.75, and 1.25 mg/kg, respectively), to mice bearing HT1376 tumors. Saline-treated mice serving as controls (**Fig. 7A**). PET imaging with [^68^Ga]AJ647 was performed at 24, 48, and 72 hours post-treatment to assess Nectin-4 engagement dynamics.

**Figure 7.**
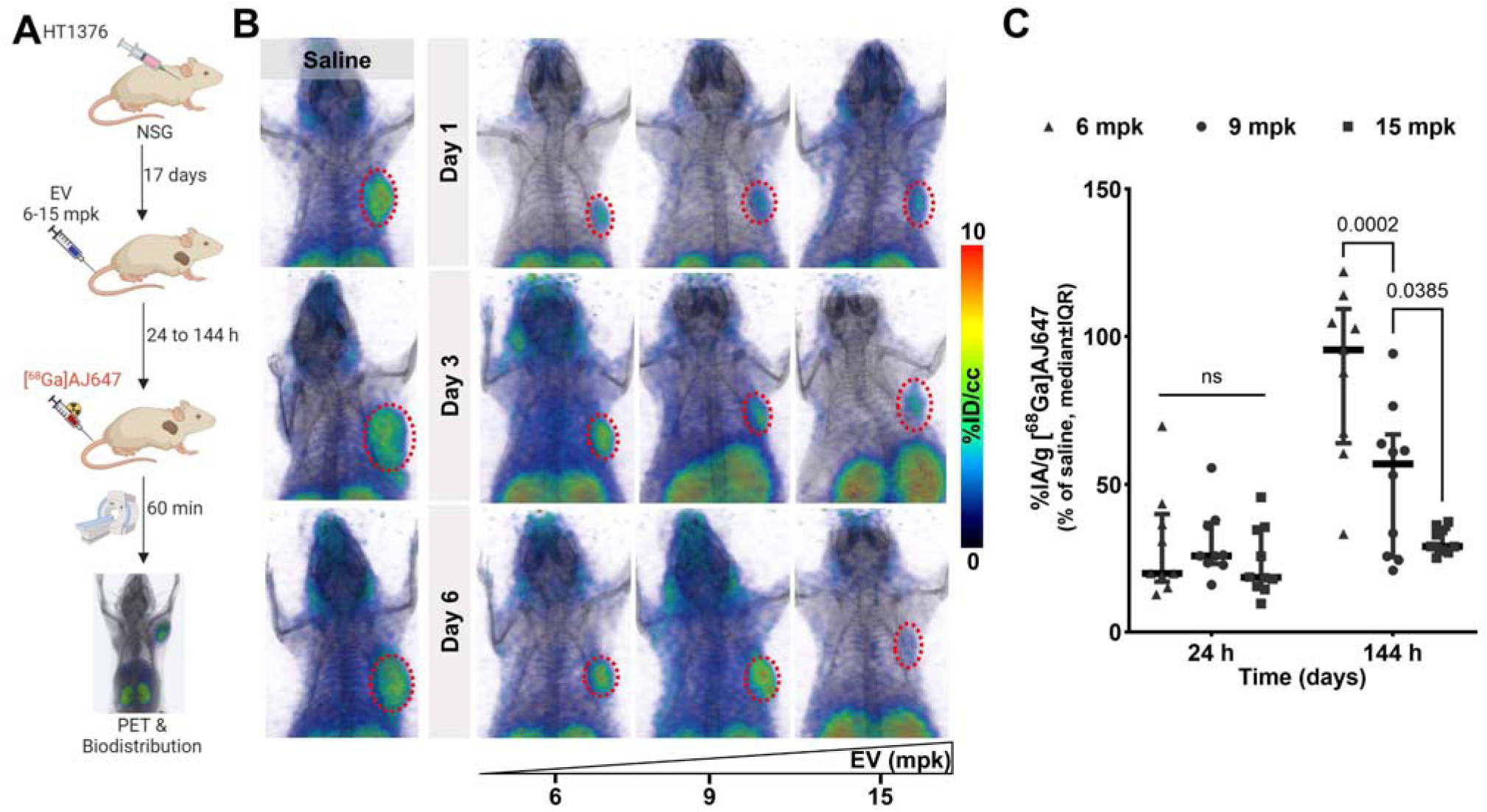
In vivo quantification of EV-Blocked Nectin-4 using [^68^Ga]AJ647-PET in HT1376 tumor xenografts. **A)** Schematic illustration of the experimental design showing how different doses of EV block Nectin-4 over various periods. **B)** Whole-body static PET-CT images of saline (left; n=2) or EV-treated mice (right; n=3) at various doses and time points, illustrating the dose- and time-dependent blocking of Nectin-4 by EV. The mice received EV at FDA-approved human doses adjusted for mouse equivalents, administered with a single dose represented on the x-axis. **C)** Quantification of EV-blocked Nectin-4 in tumor using ex vivo biodistribution data after 1 day and 6 days (n=8-9 per dose per time-point). This panel provides a comparison of the radiotracer uptake in tumors, showing a significant reduction in [^68^Ga]AJ647 binding at 6 days due to Nectin-4 blockade. Data represented as median±IQR and 2-way ANOVA used in C.

Saline-treated control mice exhibited high uptake of [^68^Ga]AJ647, indicating full availability of Nectin-4 receptors (**Fig. 7B, left**). On day 1, all EV doses effectively reduced [^68^Ga]AJ647 uptake in tumors, demonstrating full occupancy of Nectin-4 across all dose levels (**Fig. 7B, top**). By day 3, however, an increase in [^68^Ga]AJ647 uptake was observed in the tumors of mice treated with the 6 mg/kg dose, suggesting that Nectin-4 became more accessible at this lower dose as the ADC cleared from the tumor. In contrast, tumors in the 9 and 15 mg/kg groups continued to show low tracer uptake, indicating sustained target engagement (**Fig. 7B, middle**). By day 6, the dose-dependent effects became even more pronounced. The 6 mg/kg group exhibited a significant increase in [^68^Ga]AJ647 uptake, reflecting increased availability of Nectin-4, while the 9 mg/kg group showed a moderate increase in uptake, suggesting partial engagement. The 15 mg/kg group maintained minimal tracer uptake throughout, indicating persistent high-level engagement of Nectin-4 by EV (**Fig. 7B, bottom**).

Ex vivo biodistribution studies confirmed the PET imaging results by providing quantitative measurements of Nectin-4 engagement (**Fig. 7C and S10A-B**). These findings demonstrate that Nectin-4 PET imaging can effectively quantify the effects of dose and time on Nectin-4 engagement by EV, offering a valuable real-time tool for optimizing the dosing of EV and other Nectin-4–targeted therapeutics.

### Nectin-4 Dynamics and Correlation with EV Response

Further analysis of data in Fig. 7 revealed several important trends, including a dose-dependent decrease in tumor volumes at day 6 following EV treatment as well as notable variability in [^68^Ga]AJ647 uptake within tumors across different EV dose groups. Although we observed an inverse correlation between radiotracer accumulation and tumor weight (**S10C**), the relationship was weak, suggesting that radiotracer uptake may reflect tumor response to EV therapy, though other factors may contribute to this variability.

To investigate this further, we conducted experiments correlating EV dose, Nectin-4 PET measures, and response. HT1376 tumor-bearing NSG mice were randomized, treated with single doses of 6 and 15 mg/kg of EV, and tumor volumes were monitored for two weeks **(Fig. S11A)**. PET imaging using [^68^Ga]AJ647 was performed pre-treatment (day 0) and post-treatment (day 6) (**Fig. 8A**). Tumor growth inhibition was observed in both groups but was more pronounced and consistent in the 15 mg/kg group (**Fig. 8B).** The 6 mg/kg group exhibited greater variability in tumor growth response, though both groups showed a dose-dependent response.

**Figure 8.**
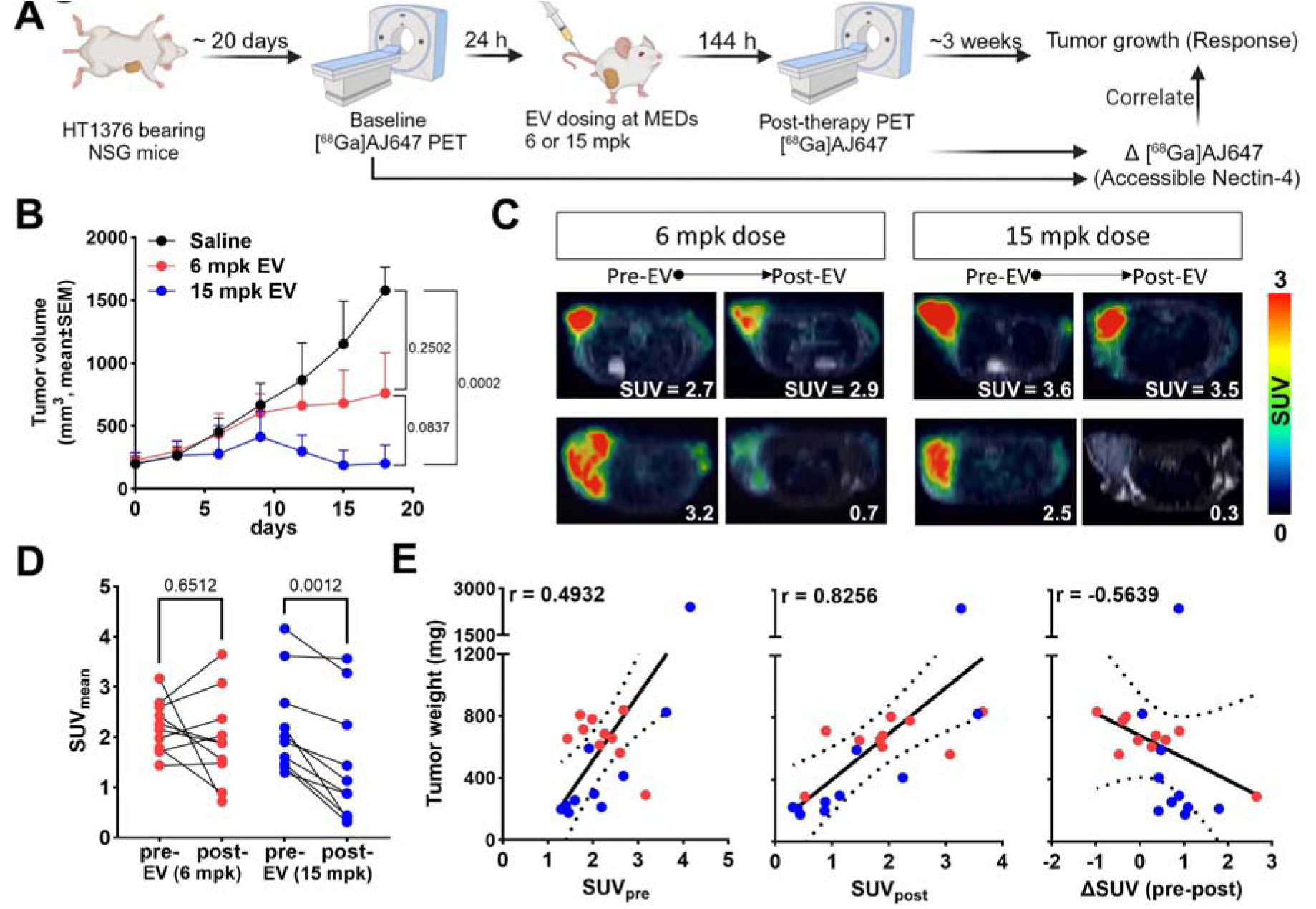
[^68^Ga]AJ647 derived Nectin-4 pharmacodynamics and correlation with response in HT1376 tumor xenograft. **A)** Schematic illustration of the experimental design showing the changes in the [^68^Ga]AJ647 tumor uptake following different doses of EV at Day 6. **B)** Tumor growth curve following single dose of EV treatment, showing growth inhibition in both EV treated groups (n=10 per group). **C)** Fused trans-axial PET-MR images showing the differences in the SUVs of [^68^Ga]AJ647 pre and post 6 days of EV treatment, indicating that EV can cause heterogenous accessible nectin-4 levels. **D)** SUVs derived for each mouse from the PET-MR imaging, showing the difference in uptake in before and after EV treatment. **E)** Correlation of tumor weights with tumor SUV before (left), after (middle) and difference (right) in before and after EV treatment. Student’s t-test used in B (unpaired) and D (paired). Spearman correlation used in E.

Analysis of PET images of those mice revealed heterogenous reduction in Nectin-4 PET signal after EV treatment in both dose groups (**Fig. 8C**). By day 6, the reduction in [^68^Ga]AJ647 uptake was more significant in the 15 mg/kg group, with 9 out of 10 mice showing a drop in uptake compared to 5 out of 10 in the 6 mg/kg group **(Fig. 8D and S11B-C)**. This suggests a stronger therapeutic effect at the higher dose. However, further analysis was needed to determine which, if any of, Nectin-4 PET measures were most relevant to predicting therapeutic response. Correlation analysis showed that pre-treatment PET measurements were only weakly associated with initial tumor volumes (r=0.3203; **Fig. S11D**) and final tumor weights (r = 0.4932; **Fig. 8E, left**). However, post-treatment standardized uptake values (SUVs) correlated more strongly with final tumor weights (r = 0.8256; **Fig. 8E, middle**), and the change in SUV from pre- to post-treatment (ΔSUV) was moderately negatively correlated with tumor weight (r = -0.5639; **Fig. 8E, right**). This suggests low uptake of [^68^Ga]AJ647 after treatment or that greater reductions in Nectin-4 PET measures may be predictive of better therapeutic outcomes.

### Target Engagement as a Predictive Biomarker for EV Therapy Response

Despite the insights gained from post-treatment SUVs, interpreting these values in isolation, without the knowledge of baseline Nectin-4 expression, poses challenges in clinical practice. This is because post-treatment SUVs can be influenced by other factors, such as baseline Nectin-4 expression variability or changes in the tumor microenvironment. Without a pre-treatment baseline, assessing the full impact of EV treatment on target engagement becomes difficult, underscoring the importance of both pre- and post-treatment PET measurements for a comprehensive evaluation of therapeutic response. To integrate these measures into analysis, we normalized ΔSUV with pre-treatment values (target engagement) and plotted SUV_post_ vs SUV_pre_, with the slope indicating the degree of target engagement (**Fig. 9A**). We found a clear relationship between target engagement and tumor response, and observed variable target engagement in both the 6 mg/kg and 9 mg/kg groups, with the higher dose group demonstrating greater target engagement.

**Figure 9.**
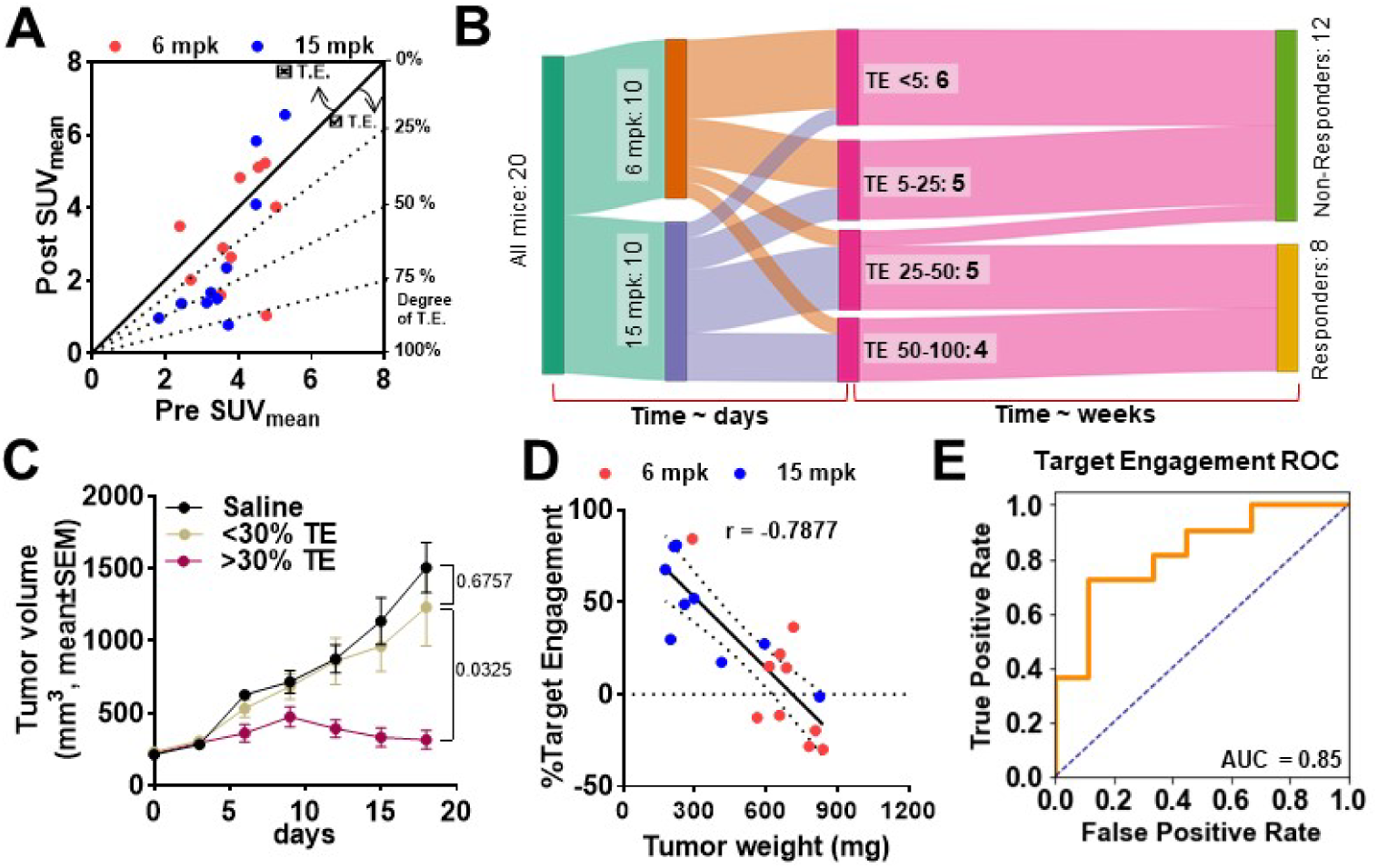
[^68^Ga]AJ647 derived EV target engagement with Nectin-4 and correlation with response. **A)** Diagonal plot of Post-treatment vs baseline SUV indicating degree of EV target engagement with nectin-4. Higher degree of nectin-4 engagement with EV is observed for points at a greater angle from diagonal. No target engagement for points lying above the diagonal line. **B)** Sanky diagram showing that irrespective of EV dose given, mice have variable target engagement, with higher target engagement leading to long-term response. **C)** Tumor growth curves based on nectin-4 engagement showing the significant tumor growth inhibition above the 25% target engagement. **D)** correlation of tumor weights with [^68^Ga]AJ647-derived % Target Engagement showing a strong negative correlation. **E)** ROC curved for target engagement shows [^68^Ga]AJ647 is a sensitive imaging tool in predicting response and dose optimization. Student’s t-test used in C (unpaired) and D (paired). Pearson correlation used in E.

Considering the variability observed in dose groups, we further sought to delineate the relationship between dose, target engagement and response. A Sankey diagram was constructed to illustrate the relationship between dose, target engagement, and response (**Fig. 9B**). We defined responders based on mRECIST criteria for mice, distinguishing between Response 0 (progressive disease) and Response 1 (stable disease or better). The flow of data showed that both dose groups exhibited variable target engagement and mice with low target engagement led to poor therapeutic outcome, regardless of the dose administered. Notably, nearly 33% of the mice showed less than 30% target engagement, while 50% exhibited less than 50% target engagement, highlighting a significant variability in response. Re-plotting the tumor volume data based on target engagement revealed that mice with over 30% target engagement exhibited significantly better responses to EV, regardless of dose (**Fig. 9C**). This re-plotting also resulted in a clearer separation of tumor growth curves compared to those plotted by dose alone (Fig. 8D). Moreover, a strong negative correlation was observed between target engagement and final tumor weight (r = -0.7877) (**Fig. 9D**), indicating that level of target engagement is predictive of therapeutic outcome.

Further analysis explored the predictive value of PET imaging-derived biomarkers, particularly target engagement, for evaluating therapeutic response. To quantitatively assess the predictive power of target engagement as a biomarker of response, we employed receiver operating characteristic (ROC) curve analysis. ROC curves provide a statistical tool for evaluating the sensitivity (true positive rate) and specificity (true negative rate) of a biomarker across varying threshold levels, enabling the identification of the optimal cutoff for distinguishing between responders and non-responders to therapy.

The ROC curves based on and pre- and post-treatment SUV values alone demonstrated poor predictive power, as indicated by area under the curve (AUC) of less than 0.5 showing poor discriminatory power of using any parameter alone **(Fig. S12A-B)**. ROC Curve of pre-post change had an area of 0.75 showing superior predictability of response **(Fig. S12C)**. In contrast, the ROC curve derived from target engagement values (ΔSUV normalized to pre-treatment SUVs) exhibited superior predictive accuracy with an area under the curve (AUC) of 0.87 (**Fig. 9E**). This stronger predictive performance indicates target engagement as a more reliable classifier for assessing therapeutic outcomes compared to pre- and post-treatment SUVs. Moreover, the ROC analysis identified a ∼30% target engagement threshold as a significant predictor of negative therapeutic outcomes **(Fig. S12D)**. Mice exhibiting less than 30% target engagement had significantly higher final tumor volume (759.4+-455 vs. 324.9+-194.6; p=0.0291) were more likely not to respond to EV treatment (0/12 responders; 0%), while those exhibiting higher target engagement may derive benefit irrespective of dose they received (7/8 responders; 87.5%). This finding suggests that patients or tumor models showing low levels of target engagement are more likely to experience poor therapeutic outcomes and may require alternate dosing or treatment strategies.

## DISCUSSION

This study successfully demonstrates the utility of Nectin-4-targeted PET imaging as a non-invasive, real-time tool for optimizing dose-exposure-response relationships in EV therapy for UC. By leveraging the specificity of [^68^Ga]AJ647 for Nectin-4, we demonstrated the ability to dynamically quantify target engagement across a range of preclinical models, bridging critical gaps in current ADC development strategies and enabling a deeper understanding of drug-tumor interactions. Importantly, we observed a strong correlation between PET-derived measures of target engagement and therapeutic outcomes. Mice exhibiting low Nectin-4 engagement post-treatment showed reduced tumor growth inhibition, regardless of dose, suggesting that target engagement itself may serve as a more reliable early biomarker of therapeutic efficacy than dose or Nectin-4 expression alone. These findings underscore the importance of integrating imaging-based biomarkers into drug development pipelines, particularly for complex therapeutics like ADCs, which have narrow therapeutic windows.

In vitro and in vivo competition studies confirmed that [^68^Ga]AJ647 binds specifically to the same epitope on Nectin-4 as EV, allowing for accurate measurement of accessible Nectin-4 levels. PET imaging quantified dose-dependent variations in target engagement, demonstrating sustained target saturation at higher EV doses while highlighting the consequences of suboptimal dosing, which led to incomplete engagement and reduced therapeutic efficacy. These insights are particularly valuable given the challenges of dose optimization in novel therapeutics. Since 2011, nine novel drugs have been granted accelerated approval based on early clinical endpoint effects but later withdrawn from the market, primarily due to failures in demonstrating improved efficacy or safety (19). Approximately 65% of these withdrawals stem from inadequate dosage optimization, which likely contributed to unfavorable benefit-risk profiles (19). Optimized dosing regimens have facilitated the reintroduction of two such drugs, including the ADC emtuzumab ozogamicin for CD33-positive acute myeloid leukemia (AML)(5), underscoring the critical need for refined dosing strategies. Our findings suggest that integrating non-invasive PET imaging into early clinical trials can address these challenges by providing complementary information on target engagement and dose-response dynamics.

This study also establishes target engagement as a predictive biomarker for therapeutic response. Using ΔSUV normalized to pre-treatment values, we identified a strong correlation between target engagement levels and tumor growth inhibition. Importantly, ROC analysis revealed that a target engagement threshold of 32% distinguished responders from non-responders, providing a quantitative benchmark for future studies. This finding aligns with emerging trends in ADC development, where integrating predictive biomarkers can enhance therapeutic precision. Biomarker use has been steadily increasing in oncology drug approvals, particularly in refining patient selection and guiding therapeutic use (20). For example, HER2-directed ADCs rely on biomarker testing to identify HER2-overexpressing tumors, with clinical responses closely linked to expression status. (21). In the case of anti–Nectin-4 EV, an IHC assay for Nectin-4 was used as a companion diagnostic, but this requirement was withdrawn early in development. However, subsequent studies revealed substantial heterogeneity in Nectin-4 expression, particularly during metastatic progression. Importantly, no significant difference in Nectin-4 levels was observed between responders and non-responders, suggesting that expression alone is insufficient to predict therapeutic outcomes. This observation underscores the importance of assessing target engagement—the actual interaction between EV and Nectin-4 within tumors. Moreover, emerging data indicates that membranous Nectin-4 expression frequently decreases during metastatic spread and correlates with EV response in patients with mUC (22). Although Nectin-4 amplification correlate with significantly higher membranous Nectin-4 protein expression and predict superior therapeutic outcomes, they are observed in only 26% of EV-treated metastatic urothelial cancer (mUC) patients (23). Given the variability in response to EV and the importance of Nectin-4 expression to response (22–24), a better understanding of the molecular basis for EV responses is crucial to improve the rational use of this effective drug for patients with mUC and to optimize its ongoing clinical development in earlier UC stages. Recent developments in Nectin-4 targeted imaging shows that PET can potentially be used to capture the heterogeneity in Nectin-4 expression within and across patients (13). Although our findings confirm the prognostic value of Nectin-4 expression, they also demonstrate that target engagement measures derived from PET imaging offer a stronger predictor of therapeutic response. The predictive potential of these early non-invasive assessments highlight the potential to refine EV dosing strategies and link noninvasive molecular visualization with therapeutic outcomes. Beyond their role in dose optimization, such biomarkers can support PK and PD assessments when changes occur in formulation, dosing regimen, or administration route, particularly in combination therapies. Our study demonstrates that early, non-invasive measures of target engagement can clarify the thresholds necessary for achieving therapeutic efficacy.

Real-time assessment of tumor heterogeneity remains a critical challenge in optimizing targeted therapies. Despite the importance of sensitive assays for real-time insights into drug-tumor interactions, such tools remain limited (6,25). Model-based approaches that integrate nonclinical and clinical data—such as PK and PD parameters—can facilitate efficacy and safety predictions(26). However, these models often fail to fully account for the dynamic complexities of tumor heterogeneity and intrinsic factors that affect drug PK/PD, especially in real-time. This is particularly true for macromolecular drugs like ADCs, where systemic exposure measurements may not adequately reflect tumor-specific drug distribution. The clinical relevance of our findings is further emphasized by comparisons to existing systemic exposure data from EV trials. In EV-301 and EV-201 trials, the average systemic exposure (C_avg_) was not a statistically significant predictor of overall survival (OS) or best overall response (BOR), though trends indicated that higher exposure quartiles correlated with response rates of 55–64% compared to 32% in the lowest quartile (27,28). These findings suggest that blood-based exposure measurements may not accurately reflect tumor-specific drug behavior. By contrast, PET imaging provided direct, tumor-specific insights into Nectin-4 engagement, enabling a more precise understanding of dose-exposure-response relationships.

Nectin-4-targeted imaging also addresses the challenge of tumor heterogeneity in UC, where variable Nectin-4 expression impacts therapeutic outcomes. While baseline Nectin-4 expression has prognostic value, our data indicate that PET-derived engagement measures are more robust predictors of response. For instance, tumors with high baseline Nectin-4 expression, such as HT1376 xenografts, exhibited greater target saturation and superior growth inhibition compared to tumors with lower expression. However, even within high-expression models, variability in engagement was observed, suggesting that PET imaging could guide personalized dosing strategies. These findings highlight the potential for Nectin-4 PET imaging to complement other biomarkers and refine ADC development. Integration with genomic or transcriptomic profiling could address interpatient variability in Nectin-4 expression and enhance response prediction. Longitudinal imaging studies could monitor dynamic changes in engagement, particularly during combination therapies or disease progression, enabling adaptive dosing strategies. Additionally, PET imaging could be employed to optimize dosing in earlier-stage UC or in patient populations with heterogeneous tumor profiles, expanding the therapeutic potential of EV.

Despite its promise, this study has limitations. While PET provides robust measures of antibody delivery and target engagement, assessing the contribution of the cytotoxic payload to therapeutic outcomes remains challenging. This complexity is compounded in ADCs like EV, where the drug-to-antibody ratio (DAR) declines over time, introducing variability in the conjugated payload’s behavior. Although both ADC and conjugated payload (acMMAE) are used as surrogates for free MMAE—the primary driver of efficacy and safety—time-dependent changes in DAR and other factors, such as internalization rates and target expression, can affect their predictive accuracy. Nonetheless, ADC exposure still correlates well with efficacy and safety endpoints in clinical contexts, suggesting that its variability may have a limited impact on overall outcomes (28–31). Future studies should focus on integrating PET-derived lesion-specific target engagement data with intracellular drug activity and conjugated payload measurements to better model therapeutic efficacy.

In summary, this study provides compelling evidence that early assessment of drug-target interactions in real-time with PET serves as a valuable tool for optimizing ADC therapy. By quantifying real-time target engagement, it addresses key limitations of conventional dose-finding paradigms and offers a pathway to enhance therapeutic precision. As ADCs and other targeted therapies continue to evolve, integrating imaging measures, like Nectin-4 PET for EV, could redefine dose selection strategies, improve patient outcomes, and accelerate the development of next-generation cancer treatments.

## MATERIALS AND METHODS

### Chemicals

NOTA-NHS ester was purchased from CheMatech, and TCEP, TIPS, HOBT, and HBTU were obtained from Chem-impex. All Fmoc protected amino acids and Fmoc-PEG_12_-COOH were purchased from Ambeed, while DIPEA, TFA, DODT, TATA, and DMF were obtained from Sigma-Aldrich. All other chemicals were purchased from Sigma-Aldrich or Fisher Scientific.

### Cell culture reagents and antibodies

All cell culture reagents were purchased from Invitrogen (Grand Island, NY). Enfortumab vedotin (EV; Nectin-4 ADC) was purchased from Johns Hopkins School of Medicine Pharmacy and used as is.

### Synthesis of AJ632

AJ632 was chemically synthesized using Liberty Blue CEM automatic peptide synthesizer employing Fmoc based solid-phase peptide synthesis on Rink Amide resin in a 0.1 mmol scale (**Fig S1**). Coupling reaction was carried out using Oxyma (0.5 mmol), DIC (1 mmol) and Fmoc-AA-OH (0.5 mmol) in DMF with microwave assisted reaction for 2 min. Fmoc group was deprotected using 20% piperidine in DMF (3 mL) for 1 min with microwave assistance. Next, PEG linker was incorporated by using HBTU (0.5 mmol), HOBT (0.5 mmol), DIPEA (0.5 mmol) and Fmoc-NH-PEG_12_-COOH (0.5 mmol) in DMF at room temperature for 1.5 h and then Fmoc group was deprotected using 20% piperidine (4 mL) in DMF for 40 min at room temperature. Once the sequence was completed on the resin, the peptidyl-resin was treated with 4 mL of cleavage cocktail (TFA:TIPS:DODT:H_2_O; 92.5:2.5:2.5:2.5) for 4 h at room temperature. The cleaved reaction mixture was precipitated with diethyl ether to obtain the linear peptide as a white solid. For the cyclization, the linear peptide (102 mg, 0.043 mmol) was dissolved in 100 mL of water:acetonitrile (1:1) and treated with an aqueous solution of tris(2-carboxyethyl)phosphine (TCEP) (105 mg, 0.37 mmol) followed by Et_3_N (400 µL, 2.9 mmol). 1,3,5-Triacryloylhexahydro-1,3,5-triazine (TATA) (30 mg, 0.12 mmol) was dissolved in 2 mL of acetonitrile and added slowly to the reaction mixture over an hour. The reaction mixture was stirred at room temperature for 48 h and quenched with TFA. The volatiles were removed, and the crude mixture was purified on a reversed-phase high-performance liquid chromatography (RP-HPLC) system using a preparative C-18 Phenomenex column (5 mm, 21.5 x 250 mm Phenomenex, Torrance, CA). The HPLC condition was gradient elution starting with 5% acetonitrile: water (0.1% TFA) and reaching 50% acetonitrile: water (0.1% TFA) in 30 min, followed by isocratic run of 50% acetonitrile: water (0.1% TFA) till 40 min at a flow rate of 8 mL/min. The product AJ632 was collected at retention time (RT) ∼30.9 min. Acetonitrile was evaporated under reduced pressure and lyophilized to form an off-white powder with a 25% yield (30 mg). The sequence of the AJ632 peptide is NH_2_-PEG_12_-(Cys-Pro-Nal-Asp-Cys-Met-hArg-dAsp-Trp-Ser-Thr-Pro-Hyp-Trp-Cys)-TATA based cyclization. This peptide was characterized by MALDI-TOF-MS. The theoretical chemical formula is C_125_H_183_N_25_O_38_S_4_ with an exact mass of 2770.2 and a molecular weight of 2772.2. The theoretical MALDI-TOF-MS mass [M + H]^+^ was 2771.2; the observed MALDI-TOF-MS [M + H]^+^ was 2773.9.

### Synthesis of AJ647

To a stirred solution of AJ632 (3.3 mg, 1.2 µmol) in 200 µL of DMF in a reaction vial, NOTA-NHS ester (2.5 mg, 2.8 µmol) and DIPEA (5 µL, 25 µmol) were added and stirred at room temperature for 4 h (**Scheme S2**). DMF was evaporated using a rotary evaporator under reduced pressure and the residual product was purified on a RP-HPLC system using a semi-preparative C-18 Luna column (5 mm, 21.5 x 250 mm Phenomenex, Torrance, CA). The HPLC condition was gradient elution starting with 5% acetonitrile: water (0.1% TFA) and reaching 50% acetonitrile: water (0.1% TFA) in 30 min, followed by isocratic run of 50% acetonitrile: water (0.1% TFA) till 40 min at a flow rate of 8 mL/min. The product AJ647 was collected at RT ∼31.2 min. Acetonitrile was evaporated under reduced pressure and lyophilized to form an off-white powder with a 27% yield, which was characterized by MALDI-TOF-MS. The theoretical chemical formula of AJ647 is C_137_H_202_N_28_O_43_S_4_ with an exact mass of 3055.3 and molecular weight of 3057.5. The theoretical MALDI-TOF-MS mass [M + H]^+^ is 3056.3, and the observed ESI-MS mass [M + H]^+^ is 3058.9.

### Affinity Measurements by Surface Plasmon Resonance (SPR)

The affinity of AJ647 for human and mouse Nectin-4 recombinant proteins were evaluated by SPR. The experiments were conducted using a Biacore T200 instrument with a CM5 chip at 25 °C. The ligands used were His-Tagged human Nectin-4 (R&D systems, catalog # 2659-N4-050, 44 kDa, 0.2 mg/ml stock concentration) and mouse Nectin-4 proteins (R&D systems, catalog # 3116-N4-050, 45 kDa, 0.2 mg/ml stock concentration), which were immobilized onto the CM5 chip. AJ647 (3057.5 Da, 10 mM stock concentration) was used as the analyte, which flowed over the ligand immobilized surface. FC2 and FC4 was used as the experimental flow cell, while FC1 and FC3 served as the reference. Anti-His antibody (1 mg/ml stock concentration) was immobilized on all FCs using standard amine coupling chemistry. The immobilization running buffer used was PBS-P (20 mM phosphate buffer pH 7.4, 137 mM NaCl, 2.7 mM KCl, 0.05% v/v surfactant P20). Human Nectin-4 was captured onto FC2 at a level of ∼600 RU, with a 1:20 dilution and 10 μg/ml diluted concentration in PBS-P. Mouse Nectin-4 was captured onto FC4 at a level of 600 RU, with a 1:20 dilution and 10 μg/ml diluted concentration in PBS-P. The theoretical R_max_ values were calculated based on the captured response values and are presented in table below, assuming a 1:1 interaction mechanism. Overnight kinetics were performed for all analytes in the presence of PBS-P+1% DMSO. The flow rate of all analyte solutions was maintained at 50 μL/min. The contact and dissociation times used were 120s and 360s, respectively. Surface regeneration was achieved by injecting glycine pH 1.5 for 20 seconds, which takes away all captured ligands onto FC2 and FC4. Fresh ligands were captured at the beginning of each injection cycle. The analyte concentrations injected ranged from 300 nM down to 1.2 nM with three-fold serial dilutions, and all analytes were injected in duplicate. All association and dissociation kinetics were evaluated by 1:1 kinetics model fitting.

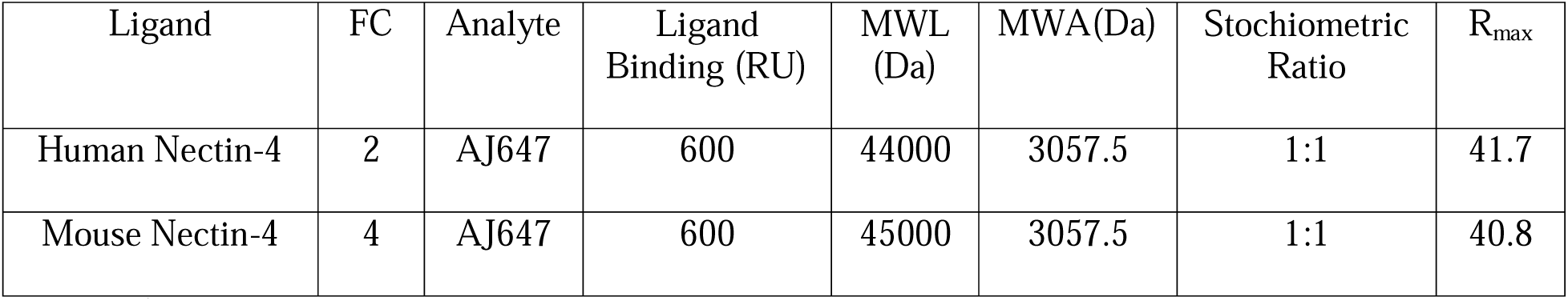

| Ligand | FC | Analyte | Ligand Binding (RU) | MWL (Da) | MWA(Da) | Stoichiometric Ratio | $R_{\max}$ |
| --- | --- | --- | --- | --- | --- | --- | --- |
| Human Nectin-4 | 2 | AJ647 | 600 | 44000 | 3057.5 | 1:1 | 41.7 |
| Mouse Nectin-4 | 4 | AJ647 | 600 | 45000 | 3057.5 | 1:1 | 40.8 |

### Synthesis of [^nat^Ga]AJ647

The synthesis of [^nat^Ga]AJ647 was carried out by adding 10 µL of aqueous 0.1M [^nat^Ga]GaCl_3_ solution and 0.6 mL of 0.1 M HCl to a stirred solution of AJ647 (0.1 mg, 0.05 µmol) in 200 µL of 1M NaOAc buffer (pH 5.0) in a reaction vial. The reaction mixture was incubated at 65 °C for 30 min and then purified on a RP-HPLC system using a semi-preparative C-18 Luna column (5 mm, 10 x 250 mm Phenomenex, Torrance, CA). The HPLC condition was gradient elution starting with 20% acetonitrile: water (0.1% formic acid) and reaching 60% acetonitrile: water (0.1% formic acid) in 20 min at a flow rate of 5 mL/min. The product [^nat^Ga] AJ647 was collected at RT ∼8.8 min. The acetonitrile was evaporated under reduced pressure and lyophilized to form an off-white powder, which was characterized by MALDI-TOF-MS. The theoretical chemical formula is C_137_H_200_GaN_28_O_43_S_4_ with an exact mass of 3122.2 and a molecular weight of 3125.2. The theoretical MALDI-TOF-MS mass [M + H]^+^ was 3123.3, and the observed ESI-MS mass [M + H]^+^ was 3124.5.

### [^68^Ga]AJ647 radiopharmaceutical preparation

The ^68^Ge/^68^Ga generator was manually eluted using 6 mL of 0.1M HCl (Ultrapure trace-metal-free) in four different fractions (2.4 mL, 1 mL, 1 mL, and 1.4 mL). To a microcentrifuge vial (1.5 mL) containing 200 μL of 1 M NaOAc buffer (pH = 5) and 40 µg of AJ647 (13 nmol), 4-6 mCi of [^68^Ga]GaCl_3_ in 0.6 mL from the second fraction was added (**Scheme S3**). The reaction mixture was incubated for 12 min at 65 °C in a temperature-controlled heating block and purified on a RP-HPLC system using a semi-preparative C-18 Luna column (5 mm, 10 x 250 mm Phenomenex, Torrance, CA). The HPLC condition was gradient elution starting with 20% acetonitrile: water (0.1% formic acid) and reaching 60% acetonitrile: water (0.1% formic acid) in 20 min at a flow rate of 5 mL/min. The radiolabeled product [^68^Ga]AJ647 was collected at RT ∼8.8 min, with decay-corrected radiochemical yield of 71.2±9.4 % (n=20). The desired radiolabeled fraction was concentrated under a stream of N_2_ at 60 °C, formulated in 10% EtOH in saline, and used for *in vitro* and *in vivo* studies. The whole radiolabeling process was completed in approximately 35 min. Quality control, stability studies, and chemical identity were also performed on the same HPLC system using the same HPLC gradient as described above.

### Determination of partition coefficient

Partition coefficients, logD (pH7.4) value, was determined according to a literature procedure (32). Briefly, a 10 µL solution of [^68^Ga]AJ647 (around 10 µCi) was added to a solution of 1-octanol (200 µL) mixed with phosphate buffered saline (PBS) (190 µL) in a 1.5 mL centrifuge tube. After the mixture was vigorously shaken and vortexed, it was centrifuged at 3000 rpm for 5 min. Aliquots (10 μL) were removed from the both phases, and the radioactivity was measured on an automated gamma counter (1282 Compugamma CS, Pharmacia/LKB Nuclear, Inc., Gaithersburg, MD). The logD was calculated as the average log ratio value of the radioactivity in the 1-octanol fraction and the PBS fraction from the three samples.

### Cell culture

All cell lines were purchased from ATCC and cultured in the recommended media in an incubator at 37°C in humid atmosphere containing 5% CO_2_. UC9, UC14, and RT112 were maintained in MEM medium. BFTC909 and SCaBER were maintained in DMEM medium. T24 cells were maintained in McCoy’s 5A medium. All cells were supplemented with 10% FBS, 1% P/S. All cell lines were authenticated using short tandem repeat profiling. All cell lines were routinely tested for mycoplasma and all experiments were performed within 10-12 passages after thawing.

### Detection of Nectin-4 expression by flow cytometry

Adherent cells were detached using enzyme free cell dissociation buffer (Thermo Fisher Scientific, Waltham, MA). Nectin-4 surface expression was evaluated by direct staining of 1×10^6^ cells in 100 μL FACS buffer (PBS with 0.1% FBS and 2 mM ethylenediaminetetraacetic acid) with phycoerythrin labeled anti-Nectin-4 antibody (clone: 337516, RnD# FAB2659P), for 30 min at 4°C. Cells were then washed and analyzed for mean fluorescence intensity (MFI) by flow cytometry.□

### *In vitro* binding assays with [^68^Ga]AJ647

*In vitro* binding of [^68^Ga]AJ647 to T24, SCaBER, RT112, UC9, UC14 and HT1376 cells was determined by incubating 1×10^6^ cells with approximately 1 μCi of [^68^Ga]AJ647 for 30 min at 4°C. After incubation, cells were washed three times with ice cold PBS containing 0.1% Tween^®^20 and counted on an automated gamma counter. All cell radioactivity uptake studies were performed in quadruplicate for each cell line and repeated three times.

### Receptor density measurements

Phycoerythrin (PE) Fluorescence Quantitation Kit (BD Biosciences #340495) containing four levels of PE/bead were used. Beads were reconstituted, with 0.5 mL of PBS containing sodium azide and 0.5% bovine serum albumin, just before use. Cells were stained for 30 min at 4°C with PE labeled anti-Nectin-4 antibody (clone: 337516, RnD# FAB2659P) and run along with the beads to estimate receptor density using flow cytometry. Calibration curve of geometric mean vs PE/bead from different bead populations was generated as per the manufacturer’s protocol. This calibration curve was used to derive receptors/cell for each cell type from their respective geometric means. Isotype controls were used to eliminate any non-specific staining. Nectin-4 receptor density measured above was then correlated with [^68^Ga]AJ647 %IA uptake measured in in vitro binding assays.

### Mouse strains and *in vivo* studies

All mouse studies were conducted through Johns Hopkins University Animal Care and Use Committee (ACUC) approved protocols. Protocol M021M175 was approved by Chair, ACUC. Xenografts were established in five-to-six-weeks old, male, non-obese, diabetic, severe-combined immunodeficient gamma (NSG) mice obtained from the Johns Hopkins University Immune Compromised Animal Core.

### Xenograft models

Mice were injected with cancer cells subcutaneously in 100 mL PBS (top right flank) for all tumor models. Following cell numbers were used: HT1376 (4 M), UC9 (3 M), UC14 (3 M), RT112 (1.5 M), T24(3 M), SCaBER (4 M). Mice with tumor volumes of 100-200 mm^3^ were used for all experiments. A minimum of 4 mice were used for all biodistribution studies. A minimum of 7 mice were used in pharmacodynamics experiment to account for greater variability.

### PET imaging of mouse xenografts

Mice with tumor volume of ∼100-150 mm^3^ were injected with ∼200 mCi (7.4 MBq) of [^68^Ga]AJ647 in 200 mL of 5% ethanol in saline intravenously and anesthetized under 2.5% isoflurane. PET-CT images were acquired at 60 minutes after radiotracer injection at 5 min/bed in an ARGUS small-animal PET/CT scanner (Sedecal, Madrid, Spain) as described (n=3 unless otherwise noted). PET-MR images were acquired on a simultaneous 7T Bruker PET-MR scanner. Dynamic imaging was conducted to evaluate pharmacokinetics of [^68^Ga]AJ647. [^68^Ga]AJ647 was injected (with help of catheter) after mice was anesthetized under 2.5% isoflurane on the equipment bed. All PET data were reconstructed using the two-dimensional ordered subsets-expectation maximization algorithm (2D-OSEM) and corrected for radioactive decay and dead time. The SUV values were calculated by finding percent of injected activity (%IA) based on a calibration factor obtained from a known radioactive quantity, then normalizing by mice weight. Amide software was used to derive SUV values by drawing region of interest to closely fit tumors. For change in accessible Nectin-4 levels following EV treatment, ΔSUV was calculated by subtracting baseline (pre-treatment) uptake from post-therapy uptake of [^68^Ga]AJ647. Target engagement was quantified by normalizing this change by initial SUV in same subject controls. Image fusion, visualization, and 3D rendering were accomplished using Amira 2020.3.1® (FEI, Hillsboro, OR).

### Immunohistochemistry

Formalin-fixed, paraffin-embedded tissue sections were soaked in xylene to remove paraffin and then rehydrated through incubations in xylene (3×5 min), 100% ethanol (2×5 min), 95% ethanol (2×5 min), 80% ethanol (2×5 min) and H_2_O (1×5 min). Antigen retrieval was performed by heating the sections in pH 8.5 ethylenediaminetetraacetic buffer in a decloaking chamber for 20 min. Endogenous peroxide activity was blocked with 3% hydrogen peroxide for 10 min followed by blocking with 10% goat serum for 1 hour, and then incubated with a primary anti-human Nectin-4 antibody (polyclonal, #ab155692, abcam) at 1:100 dilution at 4°C overnight. After washing with PBS, the secondary antibody, Signalstain Boost IHC Detection Reagent (HRP), was applied and incubated for 30 min at room temperature. The slides were washed and developed using ImmPACT DAB substrate. After washing, the slides were counterstained with Mayer’s Hematoxylin for 1 min, dehydrated using alcohol and xylene, and then cover slipped. Necrosis region was ignored when selecting fields for immunohistochemistry signal quantification. All analysis were carried out in QuPath v0.3.0.

### Ex vivo biodistribution

To validate imaging studies, *ex vivo* biodistribution studies were conducted in mice with tumors of size 100-200 mm^3^. Mice received ∼50 µCi (1.85 MBq) of [^68^Ga]AJ647 in 200 mL of 5% ethanol in saline intravenously and mice were sacrificed at 5-180 min after [^68^Ga]AJ647 injection for pharmacokinetic evaluation (n=4 per timepoint). In all other studies, 60 min time-point was chosen. Selected tissues (Blood, Skin, Muscle, Bone, Salivary glands, Tumor, Lungs, Heart, Liver, Pancreas, Spleen, Kidneys, Bladder, Testicles) were collected, weighed, counted, and their %IA/g values calculated as described previously (PMID: 37793856). For EV pharmacodynamics studies, fewer organs were harvested to accommodate more number of mice.

### In vitro competition assays

Fluorescein isothiocyanate (FITC) analog of EV was made using manufacturer recommended protocol (Thermo #46409). Briefly, 1 mg EV was incubated with FITC-NHS ester at 5:1 molar ratio overnight at 4°C at 8.0 pH 0.1M borate buffer. Excess unbound FITC-NHS ester was washed off by performing buffer exchange with 1x PBS using 3 rounds of ultra-centrifugation at 4000g for 10 minutes (3 kDa MWCO filter Amicon #UFC900308). Effect of non-radioactive AJ647 on EV binding to Nectin-4 was assessed by incubating HT1376 cells at varying concentration (5.4 µM to 0.01 nM using serial dilution) for 30 minutes at 4°C. Unbound AJ647 was washed off using 3 mL cold PBS. Cells were incubated with EV-FITC for 30 minutes at 4°C. Excess EV-FITC was washed off using 3 mL cold PBS. Mean fluorescence intensity was measured using flow cytometer and data is reported as percent of 0 nM AJ647. For effect of EV on [^68^Ga]AJ647 binding, EV was used at concentration 90 nM to 0.04 nM using serial dilutions for 45 minutes at 4°C. Unbound EV was washed off using cold PBS. Then, cells were incubated with 1 µCi [^68^Ga]AJ647 for additional 15 minutes. Excess [^68^Ga]AJ647 was washed off 3x with cold PBS. Samples were read in automated gamma counter and data is reported as percent of 1 µCi standard counts.

### Pilot in vivo experiment to study EV effect on tumor uptake of [68Ga]AJ647

Male NSG mice were implanted subcutaneously with HT1376 and SCaBER cells (4 million each) on the right and left rostral end, respectively. On day 25 after cell inoculation (average tumor volume=200±35 mm^3^), mice were randomized and treated with a single dose of either saline or 20 mpk EV ADC injected intravenously (volume not exceeding 300 µL) or with 200 µL saline (n=7/group). 24 hours following the EV dose, [^68^Ga]AJ647 PET scans (n=2/group) were acquired and ex vivo biodistribution (n=5/group) was performed using the protocols mentioned above.

### EV effect on Nectin-4 temporal pharmacodynamics

The dose-time pharmacodynamics of Nectin-4 following EV treatment was studied at doses 6, 9 and 15 mpk. Experiment involving PET imaging was performed first. Male NSG mice were inoculated with 4 million HT1376 cells at day 0. On day 23 (average tumor volume=150±30 mm^3^), mice were randomized to receive either saline, or a single dose of 6, 9 or 15 mpk EV intravenously (n=3 for each dose and n=2 for saline). Mice (n=2) from each dose group and saline were imaged on day 1, 3 and 6 following EV treatment. Once PET imaging results were analyzed, ex vivo distribution experiment was performed with n=16-18 for each dose and n=10 for saline. Treated mice (n=8-9 per dose per time-point) along with saline controls (n=5 per time-point) were sacrificed on day 1 and day 6. Biodistribution studies were performed at 60-minute after [68Ga]AJ647 injection as described above.

### Target engagement as a non-invasive metric to predict EV therapy response

Male NSG (n=35) mice were inoculated with 4 million HT1376 cells at day 0. On day 23 (average tumor volume=170±22 mm^3^), mice were randomized to receive either saline, or a single dose of 6 or 15 mpk EV intravenously (n=10 for each group). Mice (n=10) from each dose group were imaged on days 22 (pre-EV) and 29 (6 days post-EV). Saline mice (n=5) were imaged on day 25 to serve as temporal as well as treatment control. Mice body weight and tumor measurements were recorded until saline group tumors reached 1500 mm^3^ average size. Responders and non-responders were identified using modified RECIST criteria for mice published elsewhere (PMID: 26479923). Briefly, the percentage change in volume relative to baseline: % tumor volume change (ΔVol_t_) = 100% × ((V_t_ – V_initial_) / V_initial_). BestResponse was the lowest ΔVol_t_ for t ≥ 10 days. For each time t, the average ΔVol_t_ from t = 0 to t was calculated, with the BestAvgResponse defined as the lowest average ΔVol_t_ for t ≥ 10 days. Modified response criteria (mRECIST), adapted from RECIST21, were as follows: mCR (BestResponse < −95%, BestAvgResponse < −40%), mPR (BestResponse < −50%, BestAvgResponse < −20%), mSD (BestResponse < 35%, BestAvgResponse < 30%), and mPD (all others). We defined mPD as non-responders (0 response) and stable disease or better as responders (1 response). The target engagement data and response were fed into python packages ‘roc_curve’ and ‘auc’ (both from ‘sklearn.metrics’ library) to generate true positive rates (TPR) and false positive rates (FPR; ) for different target engagement thresholds. ROC curve was generated by plotting TPR (sensitivity) against FPR (1-specificity). Optimum target engagement was defined threshold at which Youden’s J Statistic (TPR-FPR) was maximum.

### Statistical analysis

All statistical analyses were performed using Prism 9.0 Software (GraphPad Software, La Jolla, CA). Unpaired Student’s t-test, one or two-way ANOVA were utilized for column, multiple column, and grouped analyses respectively. P-values < 0.05 were considered statistically significant. Correlation was done using simple linear regression without keeping constant term zero.

## Data availability statement

All data relevant to the study are included in the article, uploaded as online supplementary information. All data are available on reasonable request.

## Acknowledgements

This study was funded by NIH 1R01CA236616 (S.N.),and the 68Ge/68Ga generator was supported by NIH R01CA269235 (S.N.). Core resources (histology and imaging) were supported by NIH P30CA006973. The authors thank Ms. Xiaoju Yang and Dr. Santosh Yadav for conducting PET-CT and PET-MR studies in MRB molecular imaging service center and cancer functional imaging core. The authors also thank Drs. Aykut Üren and Purushottam Tiwari of Biacore Molecular Interaction Shared Resource (BMISR) at Georgetown University for performing SPR experiments.

