## Supplementary information for "NECTIN-4 PET FOR OPTIMIZING ENFORTUMAB VEDOTIN DOSE-RESPONSE IN UROTHELIAL CARCINOMA"

**For**

**Funding:** This study was funded by NIH 1R01CA236616 (SN) and the Ga-68 generator was supported by NIH R01CA269235. Core resources (histology and imaging) were supported by NIH P30CA006973

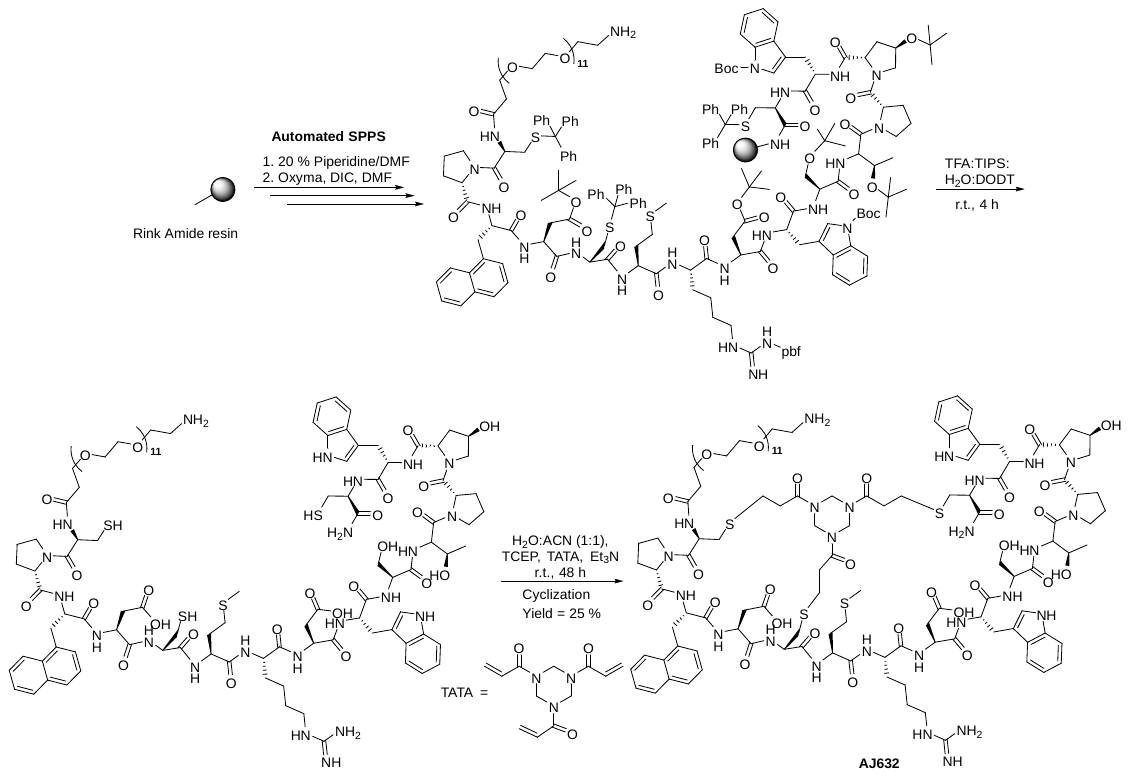

**Figure S1**. The synthesis of AJ632 was achieved through an automated solid-phase peptide synthesis (SPPS) method. First, Fmoc-protected amino acids were sequentially added to Rink amide resin via microwave-assisted coupling reactions. Next, the resulting peptidyl resin underwent treatment with a cleavage cocktail to yield the linear, deprotected peptide. Finally, the linear peptide was cyclized using TATA in the presence of triethylamine (Et3N) in a water: acetonitrile mixture, leading to the formation of AJ632.

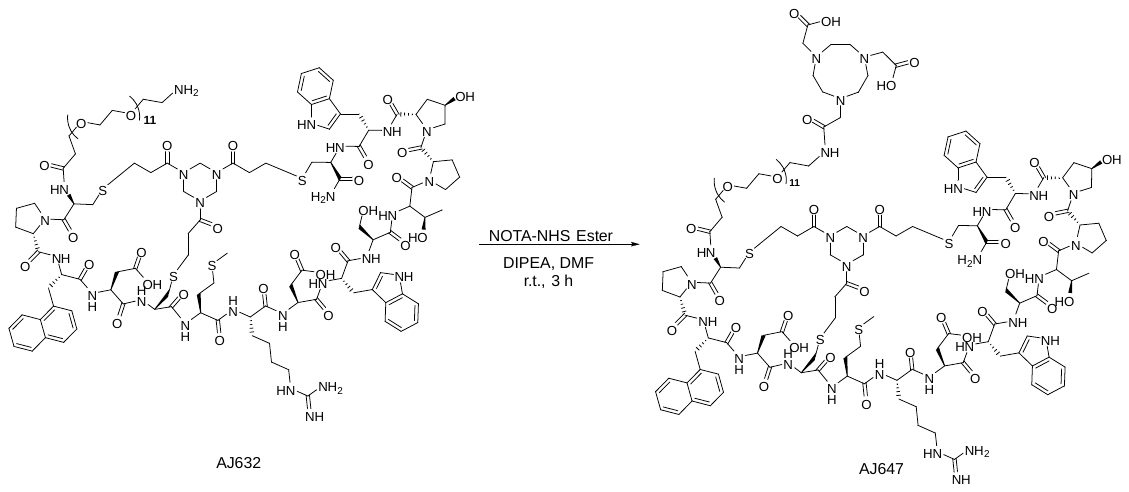

**Figure S2. Conjugation reaction of AJ632 with NOTA-NHS Ester to Obtain AJ647.** The reaction involves adding NOTA-NHS esters, a chelating agent in the AJ632, a peptide, in the presence of diisopropylethylamine (DIPEA) as a base and DMF as a solvent at room temperature for 3 hours. Following the reaction, the product, AJ647, was purified by HPLC.

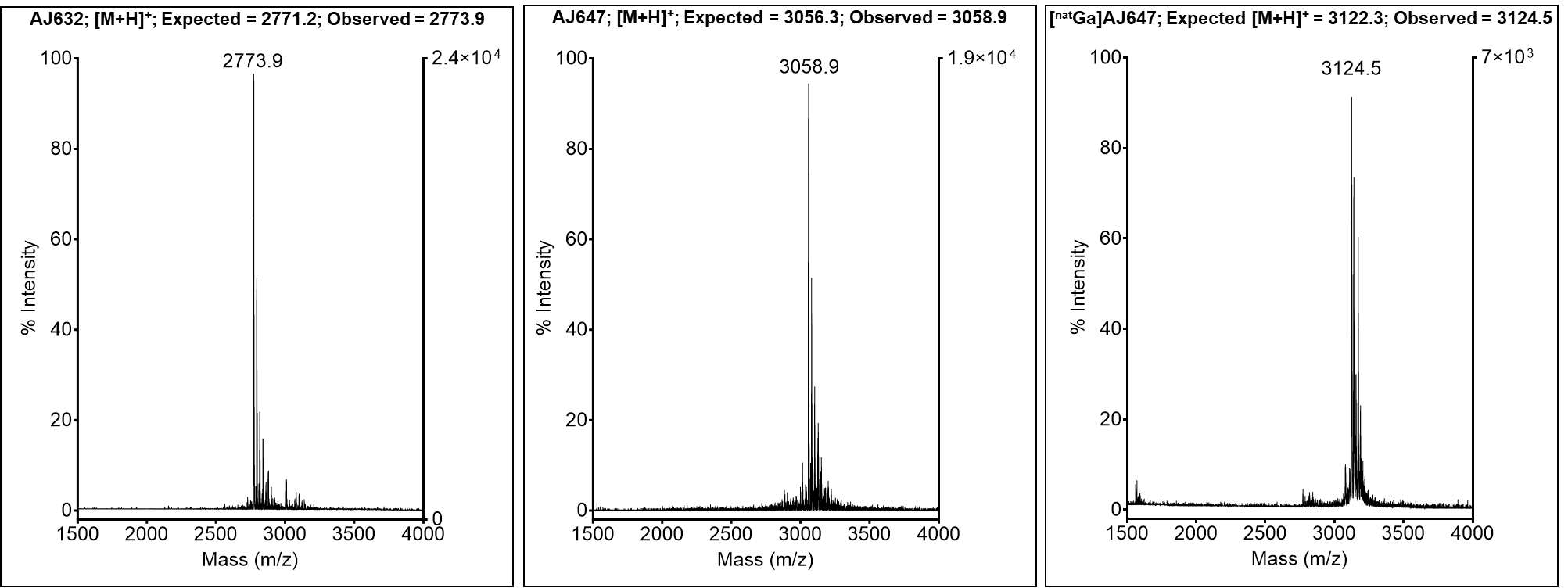

**Figure S3. Characterization of AJ632, AJ647, and [^nat^Ga]AJ647 Using MALDI-TOF Mass Spectrometry.** This figure presents the MALDI-TOF mass spectra for AJ632, AJ647, and [^nat^Ga]AJ647. Left panel shows the mass spectrum of AJ632 and expected [M+H]^+^ was 2771.2 and observed [M+H]^+^ was 2773.9. Middle panel displays the spectrum for AJ647, and expected [M+H]^+^ was 3056.3 and observed [M+H]^+^ was 3058.9, indicating successful conjugation of the NOTA chelator to the peptide. Right panel, illustrates the spectrum of [^nat^Ga]AJ647, and expected [M+H]^+^ was 3122.3 and observed [M+H]^+^ was 3124.5, indicating successful incorporation of the non-radiolabeled gallium and showing the structural chemical identity with [^68^Ga]AJ647.

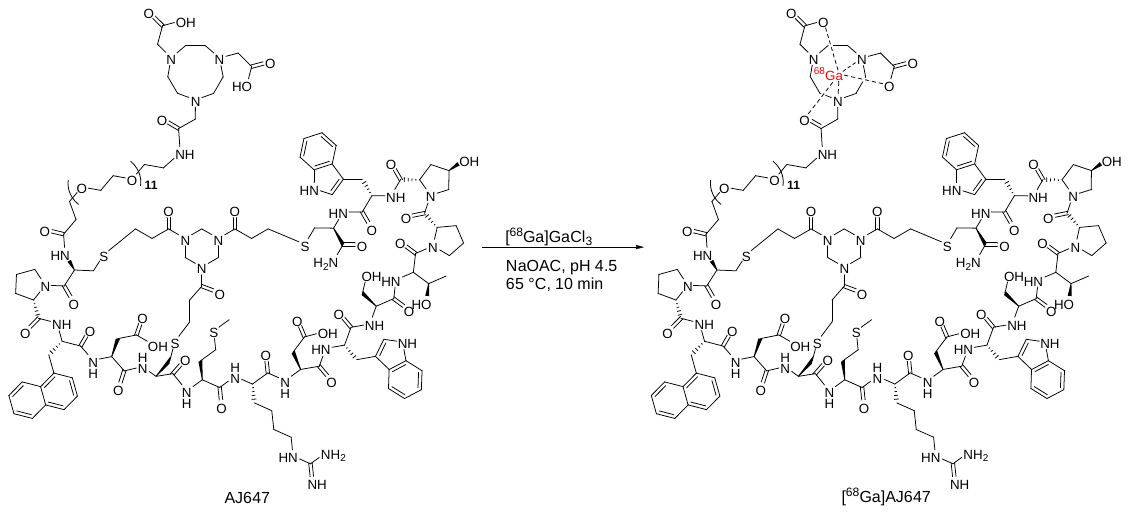

**Figure S4. Radiolabeling reaction of AJ647 with [^68^Ga]GaCl_3_**. This scheme showing the radiolabeling of AJ647 with [^68^Ga]GaCl_3_ to produce [^68^Ga]AJ647. AJ647 was reacted with [^68^Ga]GaCl_3_ in a glass vial, where the gallium-68 isotope complexes with the NOTA chelator on AJ647 under optimized heating conditions. Following the radiolabeling reaction, [^68^Ga]AJ647 was purified by HPLC to remove unreacted [^68^Ga]GaCl_3_ and other impurities, yielding the final radiolabeled product for use in in vitro and in vivo studies.

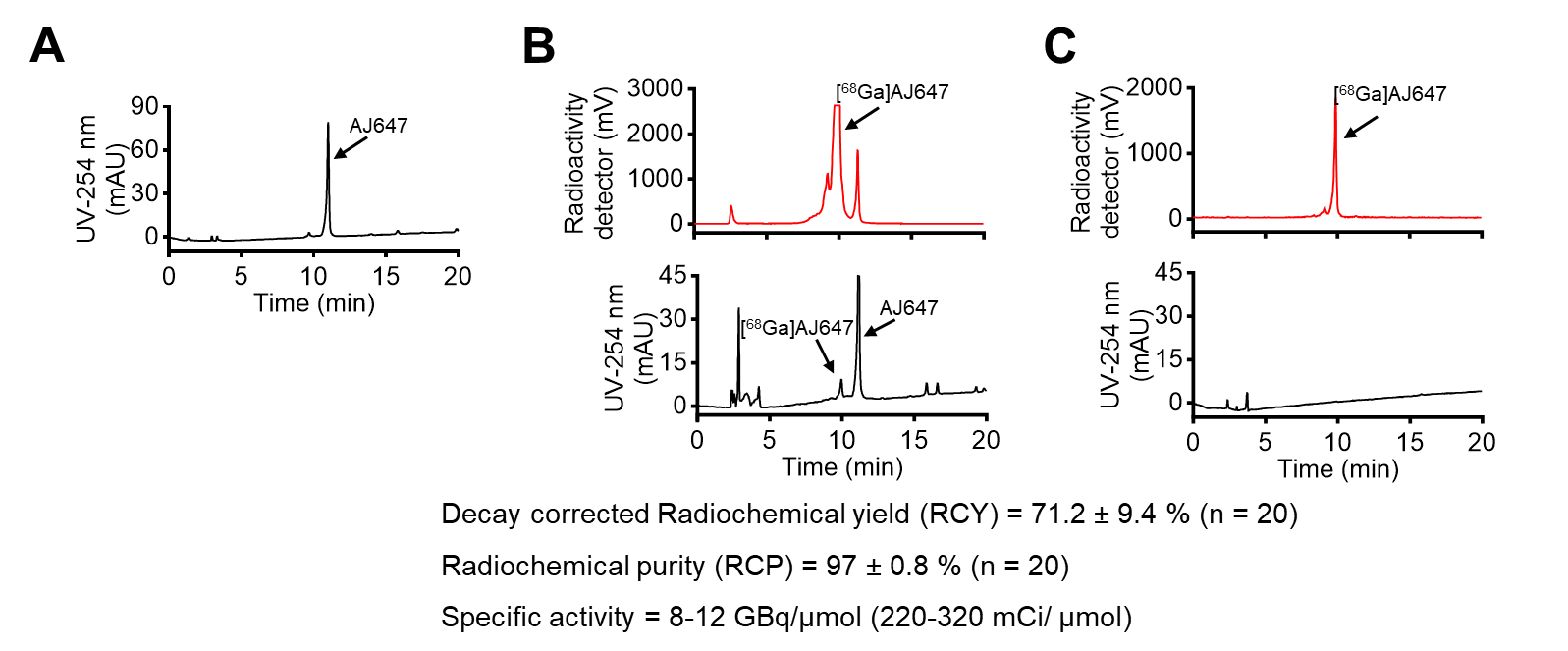

**Figure S5. Radiolabeling and Characterization of [^68^Ga]AJ647** **A)** HPLC chromatograms of AJ647 before radiolabeling. The chromatogram shows the retention time of 11.0 min and >95% of purity of the unlabeled AJ647 peptide, providing a baseline for comparison with the radiolabeled product. **B)** HPLC chromatograms of the purification process following the gallium-68 labeling reaction, showing the separation and purification of [^68^Ga]AJ647 with the decay-corrected radiochemical yield of 71.2 ± 9.4 (n = 20). **C)** HPLC chromatogram of quality control analysis of the purified radiolabeled product after formulation, showing the purity of >95 % with the molar specific activity of approximately 8-12 GBq/µmol.

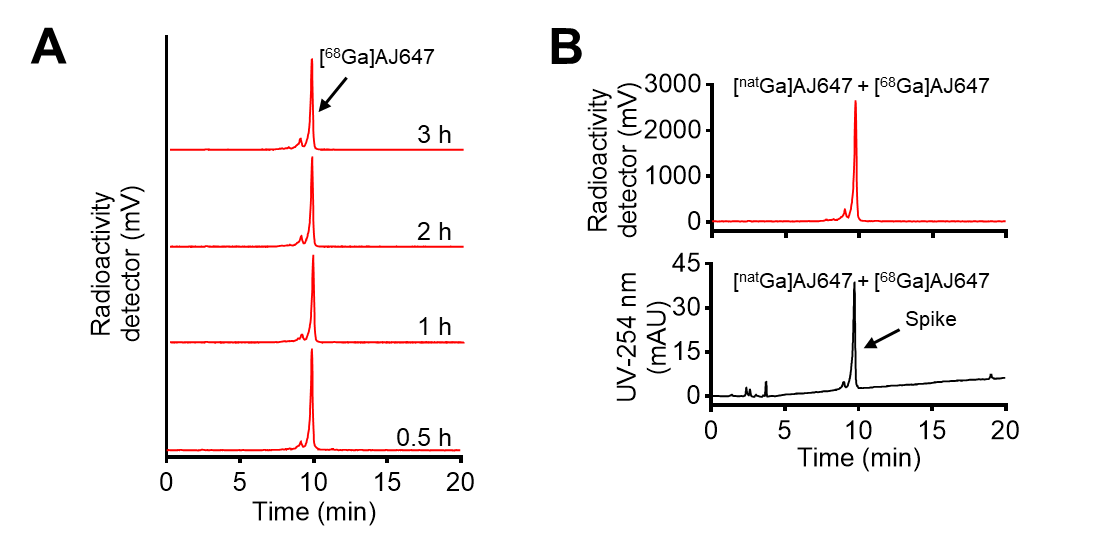

**Figure S6. Characterization of [^68^Ga]AJ647 A)** Stability of [^68^Ga]AJ647 in formulation buffer over a period of up to 3 hours. This demonstrates the radiotracer’s stability and integrity, confirming that [^68^Ga]AJ647 remains intact and reliable for imaging applications throughout this time frame. **B)** HPLC chromatograms showing the chemical identity of [^68^Ga]AJ647 with [^nat^Ga]AJ647. The chromatograms show the retention times and peak profiles of both the radiolabeled and non-radiolabeled forms of AJ647, validating the successful incorporation of gallium-68 and confirming the chemical identity of the radiolabeled product.

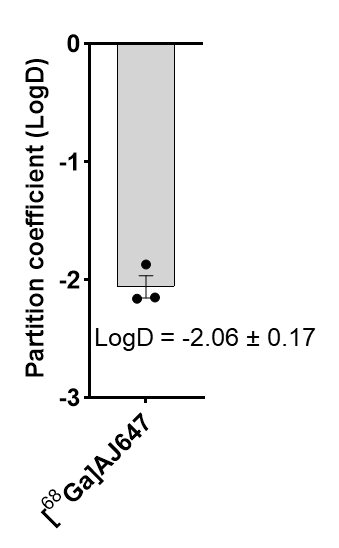

**Figure S7.** Partition coefficient of [^68^Ga]AJ647 between n-octanol and PBS. The determined Log D value was -2.06 ± 0.17, indicating high aqueous solubility of [^68^Ga]AJ647.

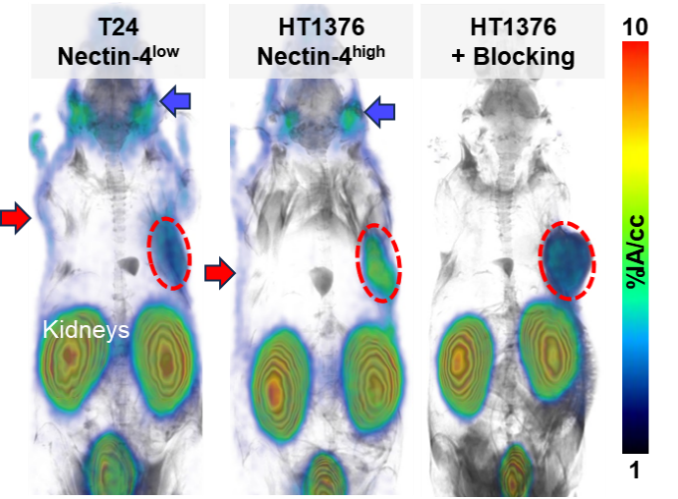

**Figure S8. [^68^Ga]AJ647 uptake in tumor xenografts, salivary glands, skin, and Nectin-4 Specificity A)** Whole-body static PET-MR imaging 60 minutes after injection of approximately 7.4 MBq (∼200 µCi) of [^68^Ga]AJ647 in NSG mice bearing T24 (low Nectin-4) and HT1376 (high Nectin-4) tumors. The images reveal high accumulation of [^68^Ga]AJ647 in the HT1376 tumor (indicated by a red circle) compared to the T24 tumor. Red arrows points to [^68^Ga]AJ647 uptake in the skin, and blue arrows indicate accumulation in the salivary glands, demonstrating Nectin-4 dependent uptake in these healthy tissues. **B)** To assess specificity, excess non-radioactive AJ647 (2 mg/kg) was pre-injected to block Nectin-4 receptors, followed by whole-body PET-MR imaging. The resulting image shows reduced radiotracer accumulation in the HT1376 tumor, confirming the specificity of [^68^Ga]AJ647 for Nectin-4.

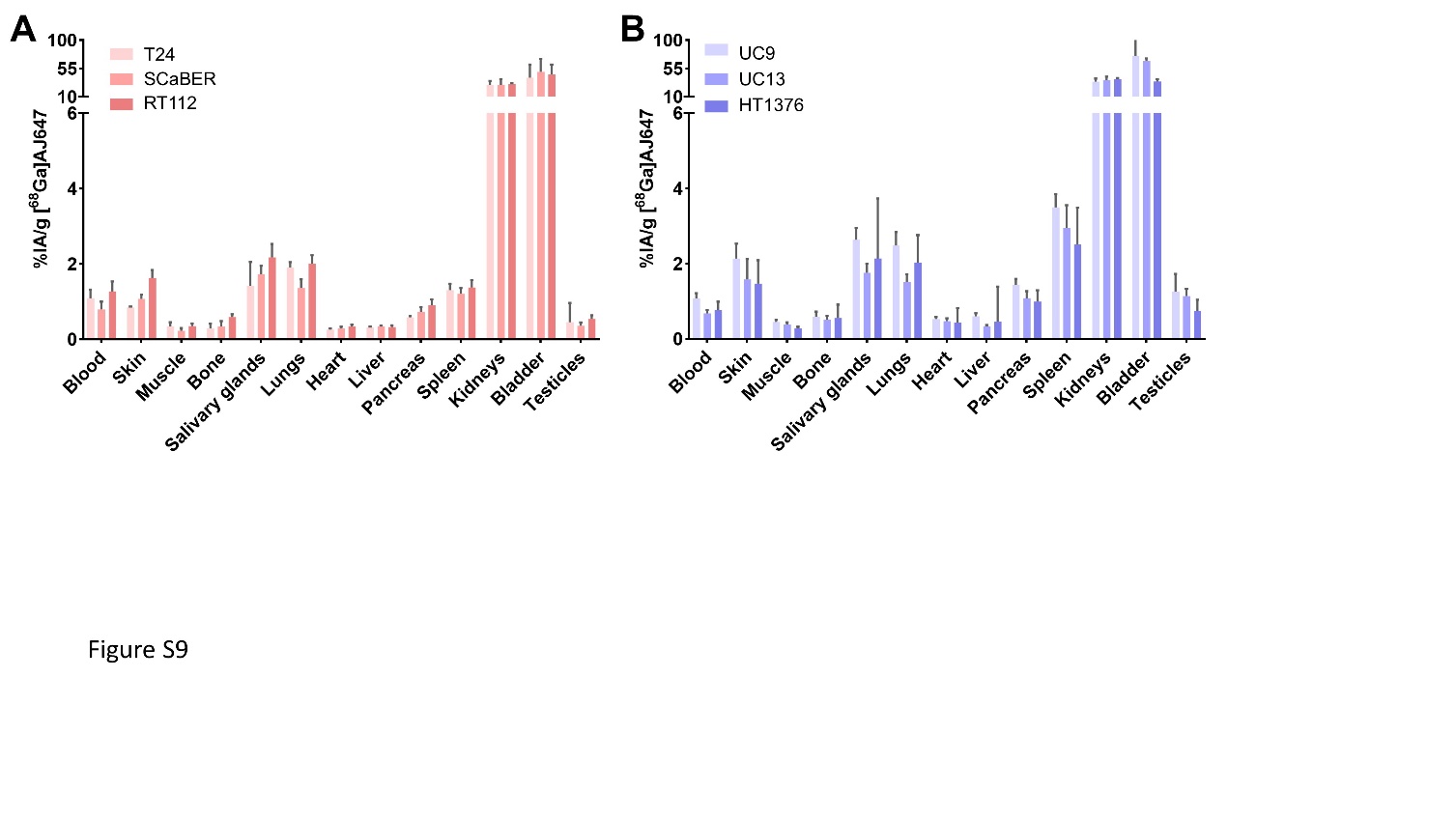

**Figure S9.** Ex vivo biodistribution of [^68^Ga]AJ647 in NSG mice bearing various bladder cancer xenografts (basal phenotypes in **A**, luminal phonotypes in **B**), as used in Figure 4. Mice were sacrificed 60 minutes after injection of approximately 740 kBq (∼20 μCi) of [^68^Ga]AJ647. The figure shows quantified radiotracer distribution in selected healthy tissues, highlighting the tissue-specific uptake and distribution patterns of [^68^Ga]AJ647. 2-way ANOVA was used to find differences between the groups and none of the differences were significant (p-value greater than 0.05).

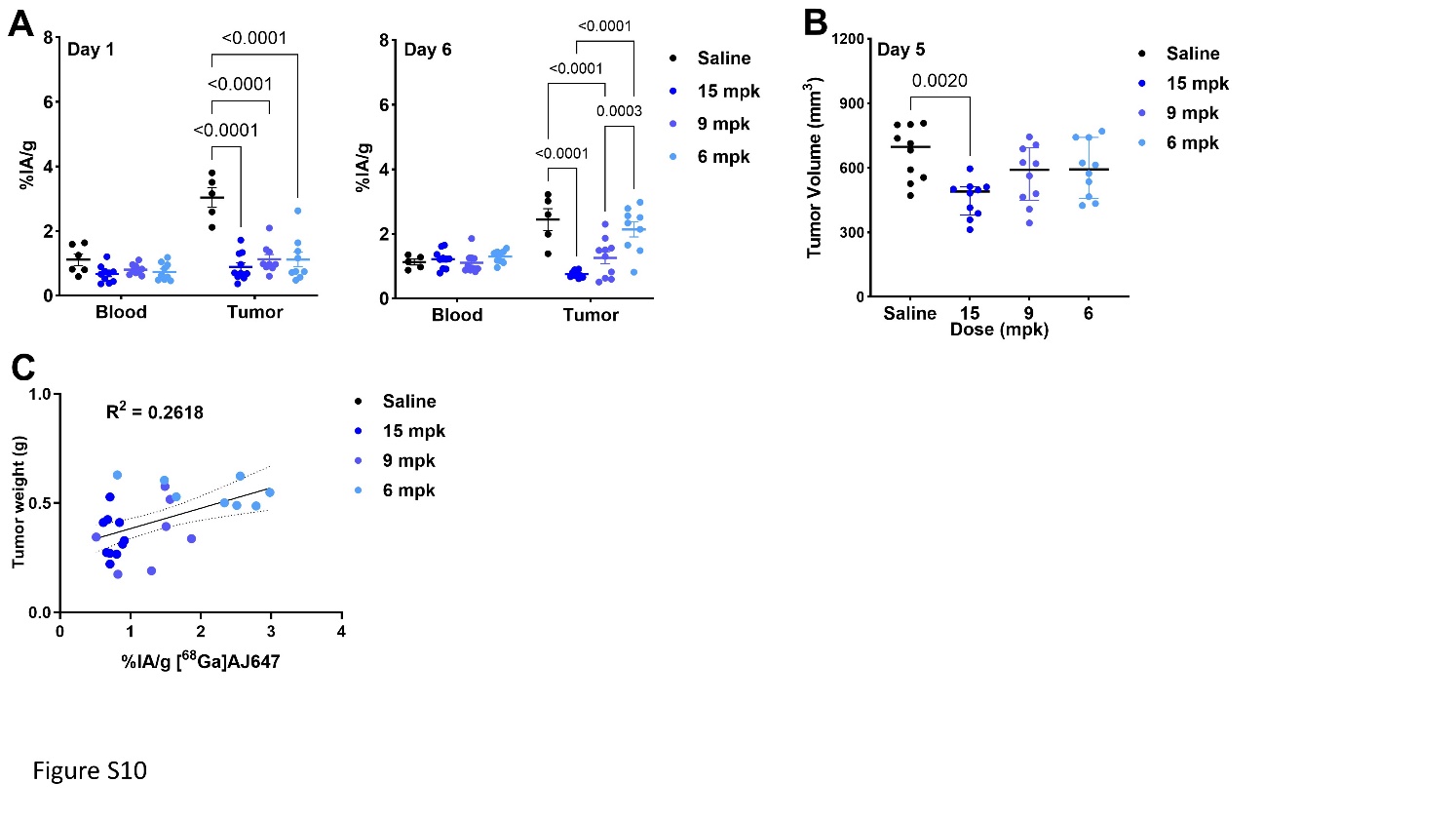

**Figure S10. Quantification of EV Nectin-4 engagement using [^68^Ga]AJ647. A)** Uptake of [^68^Ga]AJ647 in blood and tumors of mice bearing HT1376 xenografts used in Figure 6. All mice were sacrificed at 60 min after injection of ~740 kBq (~20 μCi) [^68^Ga] AJ647. 2-way ANOVA used to derive p-values. **B)** Final tumor volume of mice used in figure 6. **C)** Correlation analysis between Nectin-4 blocking and tumor weight after 6 days of EV administration, indicating a weak correlation, suggesting that [^68^Ga]AJ647 PET imaging has the potential to predict long-term tumor response to EV treatment. 1-way ANOVA used to derive p-values and pearson correlation used in C.

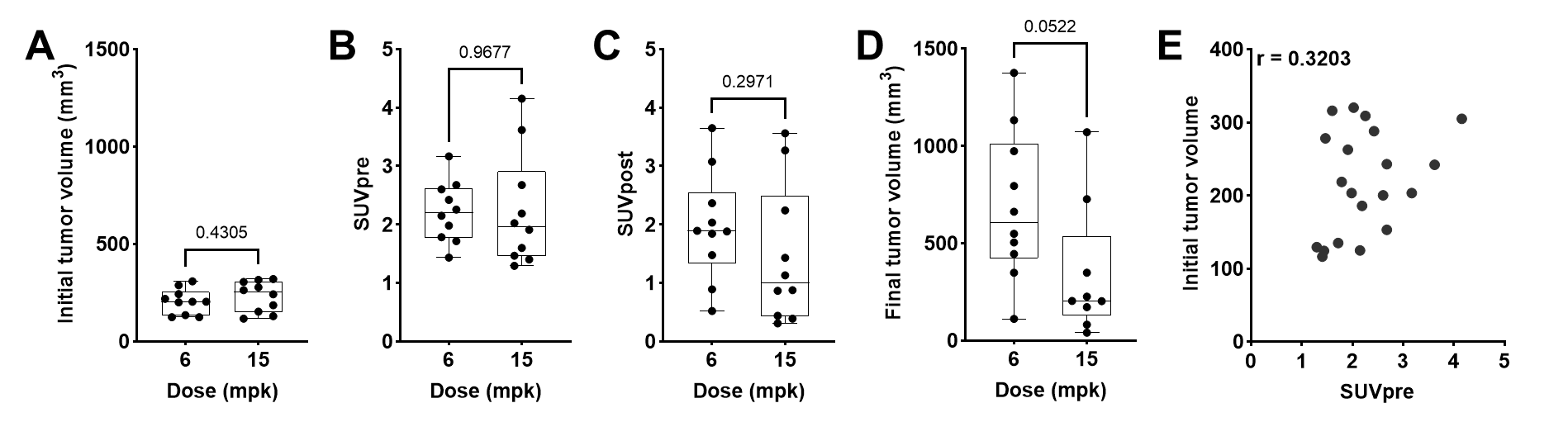

**Figure S11. Effect of dose on [^68^Ga]AJ647 uptake. A)** Mice were randomized to have similar tumor volumes before start of the treatment. [^68^Ga]AJ647 tumor **(B)** SUV_pre_ and **(C)** SUV_post_ shows subtle differences between the dose groups. **D)** Correlation between initial tumor volume and [^68^Ga]AJ647 SUV_pre_ for n=20 mice used in the study. Student’s t-test used in A-D and Pearson correlation used in E.

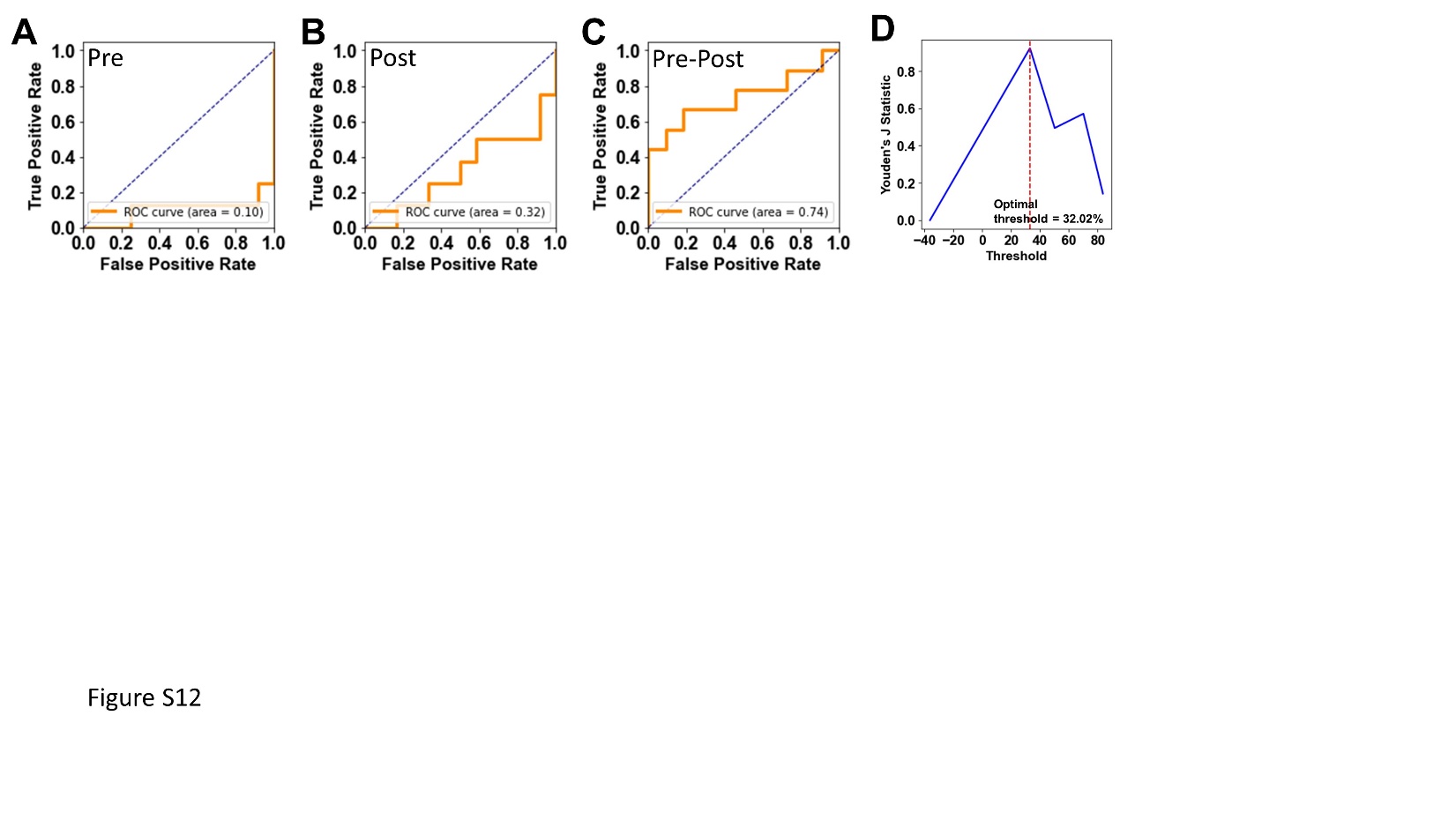

**Figure S12. Predictive analysis based on different parameters derived from [^68^Ga]AJ647 PET SUV.** ROC curves based on **A)** SUV_pre_ , **B)** SUV_post_ and **C)** ΔSUV_pre-post_ to predict long-term responders.**D)** Youden’s J statistic to find optimum threshold to differentiate responders and non-responders.
